# Striatal µ-Opioid Receptor Activation Triggers Direct-Pathway GABAergic Plasticity to Induce Negative Affect

**DOI:** 10.1101/2022.05.23.493082

**Authors:** Wei Wang, Xueyi Xie, Xiaowen Zhuang, Yufei Huang, Tao Tan, Himanshu Gangal, Zhenbo Huang, William Purvines, Xuehua Wang, Alexander Stefanov, Ruifeng Chen, Emily Yu, Michelle Hook, Yun Huang, Emmanuel Darcq, Jun Wang

## Abstract

Withdrawal from chronic opioid use often causes hypodopaminergic states and negative affect, which drives relapse. Direct-pathway medium spiny neurons (dMSNs) in the striatal patch compartment contain high levels of µ-opioid receptors (MORs). It remains unclear how chronic opioid exposure affects these MOR-expressing dMSNs and their striatopallidal and striatonigral outputs to induce negative emotions and relapse. Here, we report that MOR activation acutely suppressed GABAergic striatopallidal transmission in habenula-projecting globus pallidus neurons. Notably, repeated administrations of a MOR agonist (morphine or fentanyl) potentiated this GABAergic transmission. We also discovered that intravenous self-administration of fentanyl enhanced GABAergic striatonigral transmission and reduced the firing activity of midbrain dopaminergic neurons. Importantly, fentanyl withdrawal caused depression-like behaviors and promoted the reinstatement of fentanyl-seeking behaviors. These data suggest that chronic opioid use triggers GABAergic striatopallidal and striatonigral plasticity to induce a hypodopaminergic state, promoting negative emotions and leading to relapse.

**Highlights:**

1. Repeated administration of morphine potentiates IPSC^dMSN◊GPh^ neurotransmission.
2. Repeated administration of fentanyl potentiates IPSC^dMSN◊SNc^ neurotransmission.
3. Fentanyl withdrawal induces negative emotional states, which drive relapse.

## INTRODUCTION

Opioids are primarily used to suppress pain (<u>Reuben et al., 2015</u>). However, chronic opioid use often causes addiction and relapse (<u>Volkow et al., 2019</u>). The treatment options for opioid use disorder are limited by an incomplete understanding of the neuroplasticity mechanisms underlying opioid-induced positive and negative reinforcement. Although they bind to three types of receptors in the central nervous system, namely the mu (μ, MOR), delta, and kappa receptors, opioid-induced reward, tolerance, and dependence are primarily mediated by MORs (<u>Evans, 2004</u>; <u>George et al., 1994</u>; <u>Matthes et al., 1996</u>; <u>Volkow et al., 2019</u>; <u>Williams et al., 2013</u>). MORs are densely expressed in the midbrain, the medial habenula-interpeduncular nucleus pathway, and the striatum (<u>Bailly et al., 2020</u>; <u>Fricker et al., 2020</u>; <u>George et al., 1994</u>; <u>Georges et al., 1998</u>; <u>Jordan et al., 2000</u>). The reinforcing effects of opioids, which contribute to the onset of opioid use disorder, are mediated by activating MORs in midbrain GABAergic interneurons, causing disinhibition of dopaminergic neurons (<u>Johnson and North, 1992</u>). This disinhibition increases striatal dopamine release to reinforce drug-seeking and -taking behaviors (<u>Di Chiara and Imperato, 1988</u>; <u>Luscher and Ungless, 2006</u>).

Opioids also directly act on MORs in the striatum (<u>Jiang and North, 1992</u>), where the principal medium spiny neurons (MSNs) are located in either the direct pathway (dMSNs) or the indirect pathway (iMSNs); dMSNs project to the entopeduncular nucleus (EP; the rodent homolog of the primate internal globus pallidus or GPi) and the substantia nigra pars reticulata (SNr) and pars compacta (SNc), while iMSNs project to the external part of the globus pallidus (GPe) (<u>Bamford et al., 2018</u>; <u>Cheng et al., 2017b</u>; <u>Gerfen and Surmeier, 2011</u>; <u>Kreitzer and Malenka, 2008</u>; <u>Maia</u> <u>and Frank, 2011</u>). The striatum contains neurochemically defined sub-compartments termed patches (striosomes), surrounded by a matrix compartment (<u>Bolam et al., 1988</u>). In contrast to the matrix, the patch compartment densely expresses MORs and contains more dMSNs than iMSNs (<u>Brimblecombe and Cragg, 2017</u>; <u>Fujiyama et al., 2011</u>; <u>Gerfen, 1984</u>; <u>Graybiel and Ragsdale, 1978</u>; <u>Johnston et al., 1990</u>; <u>Prager and Plotkin, 2019</u>; <u>Smith et al., 2016a</u>). The patch and matrix compartments have different input and output connections and mediate distinct striatal processes involved in learning (<u>Crittenden and Graybiel, 2011</u>; <u>McGregor et al., 2019</u>; <u>Smith et al., 2016a</u>). Projections from patch dMSNs to the habenula-projecting globus pallidus (GPh) are engaged during action-outcomes evaluation and thus influencing reinforcement learning (<u>Stephenson-Jones</u> <u>et al., 2016</u>). The patch compartment also plays a critical role in affective and motivational processes involved in neurological disorders (<u>Crittenden and Graybiel, 2011</u>, <u>2016</u>; <u>Friedman et al.,</u> <u>2017</u>; <u>Friedman et al., 2015</u>; <u>Friedman et al., 2020</u>). A specific group of patch dMSNs encodes aversive responses and drives negative reinforcement (<u>Xiao et al., 2020</u>). Patch dMSNs are the sole source of connection between striatal activity and SNc dopaminergic neurons (<u>Crittenden et</u> <u>al., 2016</u>; <u>Fujiyama et al., 2011</u>; <u>Fujiyama et al., 2015</u>; <u>Gerfen, 1984</u>; <u>Smith et al., 2016a</u>). Withdrawal from chronic opioid exposure promotes the sensitivity to aversive stimuli, and this drives negative reinforcement (<u>Di Chiara and North, 1992</u>). This negative emotional state (hyperkatifeia), which drives relapse, is associated with hypodopaminergic activity after withdrawal from chronic opioid use (<u>Koob, 2021</u>). Therefore, the patch compartment is the direct striatal target of opioids, and it may mediate profound effects on drug reinforcement and relapse. However, it is unclear how acute and chronic opioid exposure alters the activity of patch dMSNs or their outputs to the EP and SNc.

This study found that acute MOR activity induced by an agonist (DAMGO) inhibited GABAergic striatopallidal transmission from dMSNs to GPh neurons. In contrast, repeated non-contingent administration of morphine or fentanyl potentiated this transmission. We also discovered that intravenous self-administration (IVSA) of fentanyl enhanced GABAergic transmission not only in MOR-expressing patch neurons but also in SNc dopaminergic neurons, which reduced their firing. Importantly, fentanyl withdrawal induced depression-like behavior and triggered relapse. Our data indicate that chronic opioid exposure triggers GABAergic striatopallidal and striatonigral transmission, inhibits SNc dopaminergic activity and causes negative affect. This result provides insights into the mechanism underlying opioid-induced negative emotional states, which may drive relapse.

## RESULTS

### MOR activation acutely suppresses striatopallidal GABAergic transmission in GPh neurons

GPh neurons receive GABAergic inputs from striatal patches, which mainly express MORs and predominantly contain dMSNs (<u>Stephenson-Jones et al., 2013</u>; <u>Stephenson-Jones et al., 2016</u>). Thus, we first tested whether MOR activation altered GABAergic transmission from the striatum to the EP, which contains GPh neurons. To achieve this, we prepared sagittal slices from D1- Cre;Ai167 (or Ai32) mice in which D1R-expressing neurons, including dMSNs, contained the excitatory opsin, Chrimson, and either tdTomato (tdT; Fig. 1A) or ChR2-EYFP. Optically induced inhibitory postsynaptic currents (oIPSCs) in EP neurons were completely abolished by a GABA_A_ receptor (GABA_A_R) antagonist, picrotoxin (Fig. 1B, C), confirming GABAergic transmission. We discovered that bath application of a MOR agonist, DAMGO, dose-dependently suppressed oIPSCs (Fig. 1D, E; *t*_(7)_ = 4.73, *p* < 0.01). This result suggests that MOR activation acutely inhibits GABAergic transmission from D1R-expressing neurons to the EP. In addition, bath application of DAMGO increased the paired-pulse ratios (PPRs) of oIPSC amplitudes (Fig. 1F; *t*_(6)_ = -6.94, *p* < 0.001), suggesting that MOR activation reduces presynaptic GABA release from D1R- expressing fibers to EP neurons (<u>Zucker and Regehr, 2002</u>). Moreover, bath application of a MOR antagonist, CTAP, completely abolished the DAMGO-induced suppression of oIPSCs (Fig. 1G; *F*_(2,10)_ *=* 0.07, *p* > 0.05) and increase in PPRs (Fig. 1H; *F*_(2,10)_ = 1.82, p > 0.05). These data indicate that MOR activation acutely suppresses GABAergic transmission from D1R-expressing neurons to the EP, at least in part by reducing GABA release.

**Figure 1.**
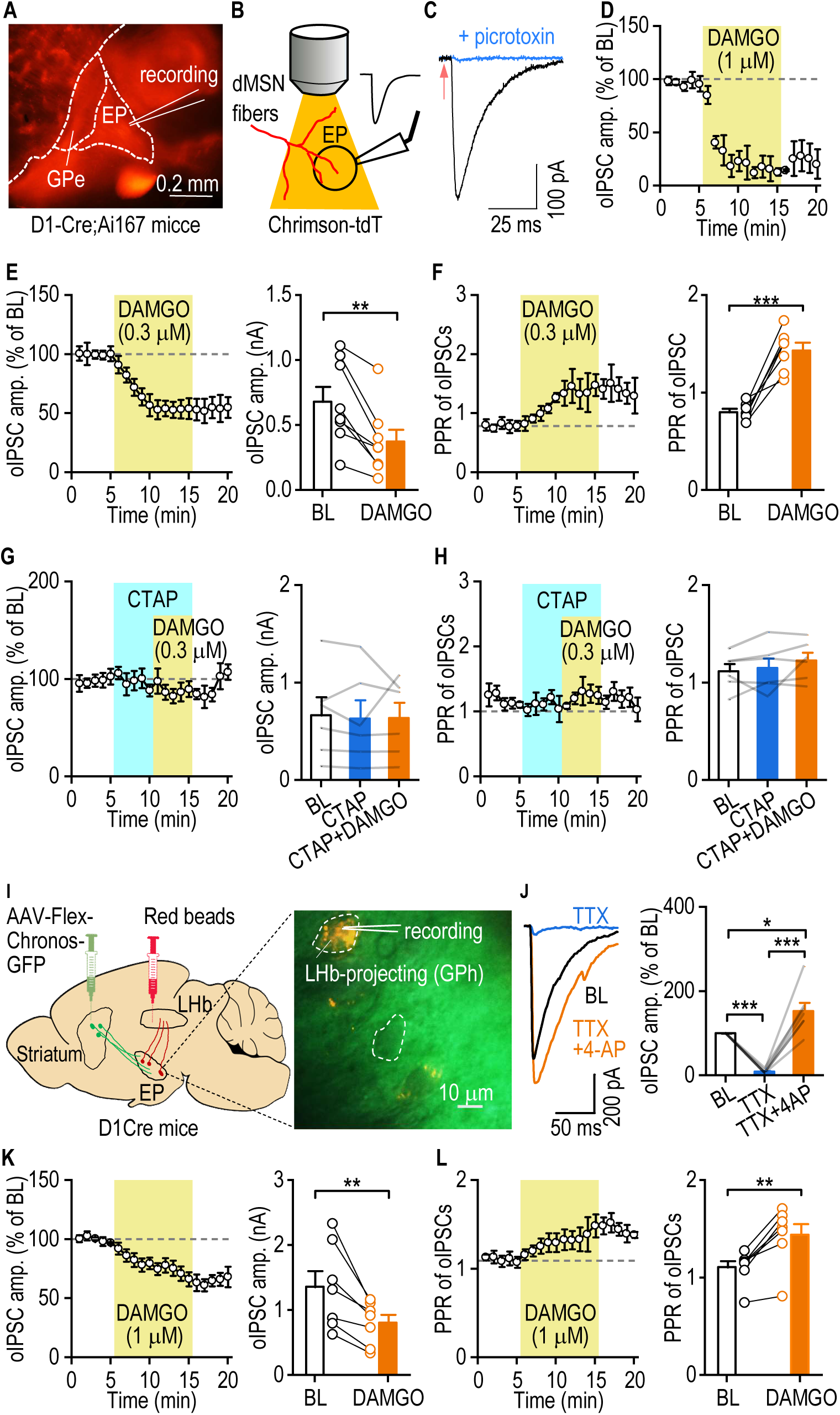
MOR activation acutely suppresses striatopallidal GABAergic transmission in GPh neurons. ***A***, Sample image of a sagittal section from a D1-Cre;Ai167 mouse showing D1R-tdTomato (tdT)- expressing fibers in the entopeduncular nucleus (EP), where whole-cell recordings were conducted. Note that this mouse expressed Chrimson-tdT in all D1R-expressing neurons and their axonal terminals in the EP. GPe, external segment of globus pallidus. ***B***, Schematic showing light stimulation of D1R^+^ Chrimson-expressing fibers and whole-cell recording in an EP neuron. EP neurons were recorded from D1-Cre;Ai167 (or D1-Cre;Ai32) mice, and light (2 ms, 590 nm for Ai167 mice or 470 nm for Ai32 mice) was delivered to selectively stimulate the D1R^+^ afferents. ***C***, Sample traces showing that optically evoked inhibitory postsynaptic currents (oIPSC) were recorded in the absence and presence of picrotoxin (100 µM). ***D, E***, A MOR agonist, DAMGO, dose-dependently (1 µM, *D*; 0.3 µM, *E*) and persistently reduced oIPSC amplitudes; \*\**p* < 0.01. BL, baseline. ***F***, DAMGO (0.3 µM) increased the oIPSC paired-pulse ratio (PPR); \*\*\**p* < 0.001. ***G, H***, A MOR antagonist, CTAP (1 μM), completely abolished the inhibitory effects of DAMGO (0.3 µM) on the oIPSC amplitude (*G*) and PPR (*H*); *p* > 0.05. ***I***, Schematic of the experimental design (left) and a sample image of recording in an LHb-projecting GPh neuron (right). AAV-Flex-Chronos-GFP and red beads were infused into the DMS and LHb of D1-Cre mice, respectively. ***J***, oIPSCs were blocked by TTX (1 μM) and recovered by adding 4-AP (500 μM); \**p* < 0.05, \*\*\**p* < 0.001. ***K, L***, DAMGO (1 µM) acutely inhibited the oIPSC amplitude (*K*) and PPR (*L*); \*\**p* < 0.01. Paired *t* test (*D*–*F*, *K*, *L*) and one-way RM ANOVA (*G*, *J*, *H*); n = 3/1 (*D*), 8/3 (*E*), 7/3 (*F*), 6/3 per group (*G*, *H*), 7/2 (*J*), 7/3 (*K*), 7/4 (*L*). D1-Cre;Ai167 (*D*), D1-Cre;Ai32 (*E*–*H*), and D1-Cre (*I–L*) mice were used.

While dMSNs express D1Rs, these receptors are also found brain-wide (<u>Hall et al., 1994</u>; <u>Lu et al., 2019</u>; <u>Mansour et al., 1992</u>; <u>Tritsch and Sabatini, 2012</u>; <u>Wei et al., 2018</u>). Next, we tested whether selective MOR activation in DMS dMSNs suppressed their outputs to GPh neurons (<u>Stephenson-Jones et al., 2016</u>), which play an important role in regulating the non-motor functions of the basal ganglia; these include action evaluation and decision-making (<u>Mollick et al., 2020</u>; <u>Stephenson-Jones et al., 2016</u>). We infused AAV-Flex-Chronos-GFP into the DMS of D1-Cre mice for selective optogenetic stimulation of dMSN outputs and infused red beads into the LHb. These beads were retrogradely transported and thus facilitated selective recording of GPh neurons (Fig. 1I). Optical stimulation induced oIPSCs in bead-labeled GPh neurons that were abolished by tetrodotoxin (TTX) and recovered by application of TTX + 4-AP (<u>Cruikshank et al., 2010</u>; <u>Petreanu</u> <u>et al., 2009</u>) (Fig. 1J; *F*_(2,12)_ = 39.9, *p* < 0.001). These data suggest that striatal dMSN outputs form monosynaptic connections with GPh neurons. This oIPSC from dMSNs to GPh neurons (oIPSC^dMSN◊GPh^) was also suppressed by bath-application of DAMGO (Fig. 1K; *t*_(6)_ = 3.76, *p* < 0.01). Moreover, DAMGO significantly increased PPRs (Fig. 1L; *t*_(6)_ = 5.45, *p* < 0.01). These data demonstrate that MOR activation acutely suppressed striatopallidal GABAergic transmission from dMSNs to GPh neurons via a presynaptic mechanism (<u>Zucker and Regehr, 2002</u>).

### Noncontingent opioid administration potentiates striatopallidal GABAergic transmission in GPh neurons

Having shown that acute MOR activation suppressed dMSN→GPh transmission, we next examined the effects of repeated MOR activation. D1-Cre mice received the same viral and bead injection depicted in Figure 1I (Fig. 2A). We first tested whether repeated injections of morphine, a widely used MOR agonist (<u>Jordan et al., 2000</u>), altered locomotor activity, which is regulated by striatal dMSNs (<u>Kravitz et al., 2010</u>). Mice received an intraperitoneal (i.p.) injection of morphine (20–100 mg/kg) once daily for 5 d, and locomotor activity was measured immediately after each injection (Fig. 2A). We found that morphine injections significantly increased the distance traveled (Fig. 2B; *F*_(1,8)_ = 83.31, *p* < 0.001) and velocity (Fig. 2C; *F*_(1,8)_ = 67.26, *p* < 0.001). Consistently, the time spent in the periphery was significantly higher in the morphine-treated group than in the saline-treated group (Supplementary Fig. 1A; *F*_(1,8)_ = 18.74, *p* < 0.01). These data indicate that i.p. morphine administrations increase locomotor activity, consistent with previous studies (<u>Hnasko et</u> <u>al., 2005</u>). Electrophysiological recordings revealed that repeated morphine injections significantly enhanced the amplitude of oIPSC^dMSN◊GPh^, as compared to saline (Fig. 2D; *F*_group (1,27)_ = 13.39, *p* < 0.01; *F*_interaction (4,108)_ = 5.85, *p* < 0.001). The oIPSC PPRs were significantly smaller in morphine-injected animals than in saline-injected animals (Fig. 2E; *t*_(23)_ = 3.12, *p* < 0.01), suggesting an increased probability of GABA release. Consistently, we observed that the frequency, but not the amplitude, of spontaneous IPSCs (sIPSCs) was significantly greater in the morphine-treated animals than in the saline group (Fig. 2F; *t*_(22)_ = - 3.3 *p* < 0.01). Together, these data suggest that repeated non-contingent morphine administration enhances dMSN→GPh GABAergic transmission and that this effect results from an increase in presynaptic dMSN GABA release.

**Figure 2.**
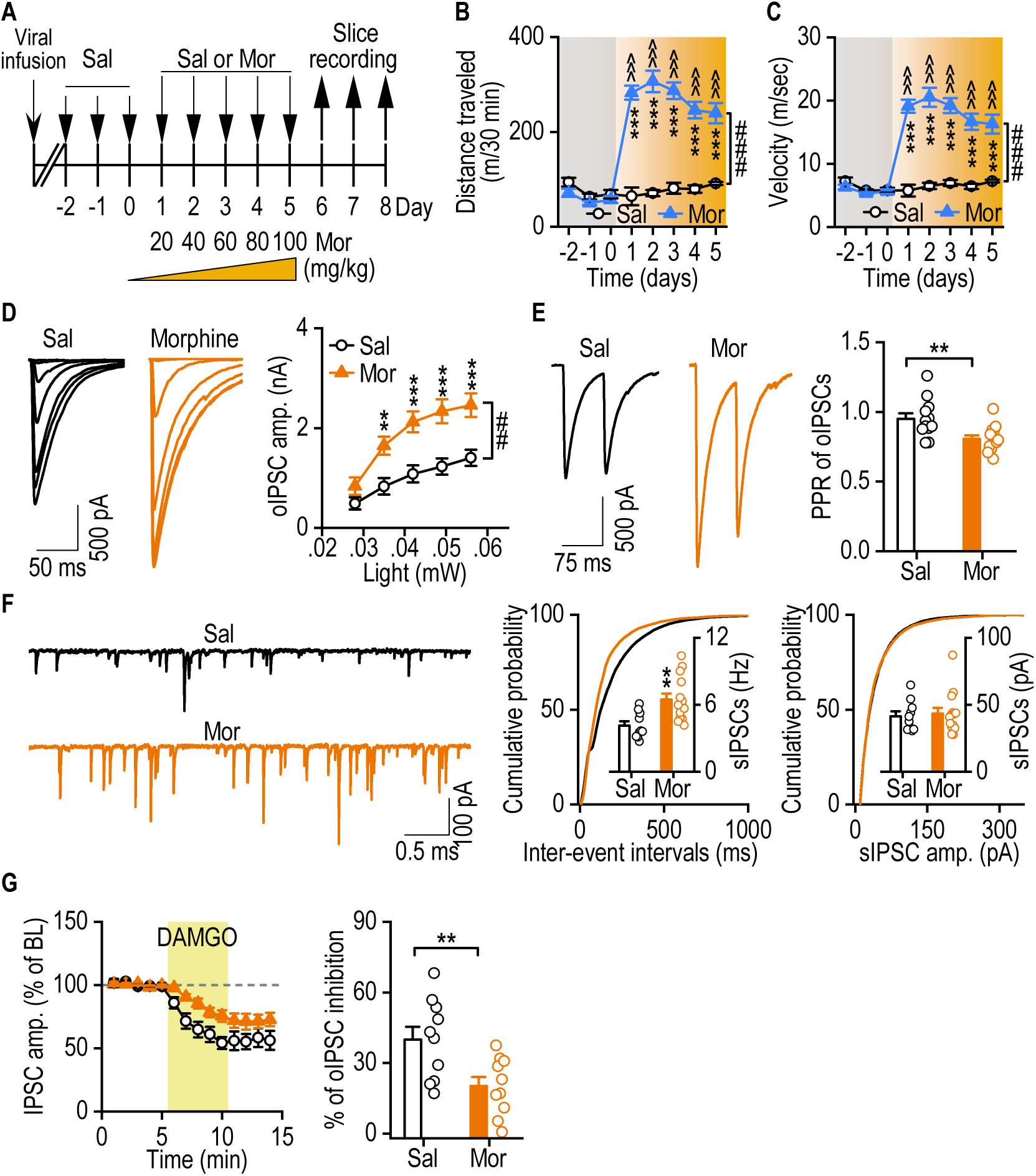
Repeated morphine administration enhances striatopallidal GABAergic transmission in GPh neurons. ***A***, Schematic showing the experimental design, with viral infusion, i.p. morphine (Mor) or saline (Sal) injections, and slice recording in D1-Cre mice. The viral infusion was the same as shown in Fig. 1I. ***B, C***, Morphine injections enhanced the distance traveled (*B*) and velocity (*C*) in locomotion tests; ^###^*p* < 0.001; \*\*\**p* < 0.001 versus the same treatment day in the saline group; ^^^*p* < 0.001 versus the last saline injection day (day 3). ***D***, Repeated morphine injections increased the amplitude of oIPSCs from dMSNs to GPh neurons; ^##^*p* < 0.01; \*\**p* < 0.01, \*\*\**p* < 0.001 versus the same stimulating intensity in the saline group. ***E***, Repeated morphine injections decreased oIPSC PPRs in GPh neurons; \*\**p* < 0.01. ***F***, Repeated morphine administrations increased the frequency, but not the amplitude, of spontaneous IPSCs; \*\**p* < 0.01. ***G***, Repeated morphine injections attenuated DAMGO (0.3 µM)-induced inhibition of oIPSCs from dMSNs to GPh neurons; \**p* < 0.001. Two-way RM ANOVA followed by Tukey *post hoc* test for *B*–*D* and unpaired *t* test for *E*–*G*; n = 4 (*B, C*, Sal), 6 (*B, C*, Mor), 14/4 (*D*, Sal),15/4 (*D*, Mor), 12/4 (*E*, Sal), 13/4 (*E*, Mor), 10/4 (*F*, Sal), 14/5 (*F*, Mor), and 10/5 (*G*, per group).

The enhanced GABAergic transmission induced by repeated morphine injections may reduce the acute effect of MOR activation. To test this possibility, we recorded oIPSCs in the presence of DAMGO. We discovered that DAMGO-induced inhibition of oIPSCs was significantly weakened in the morphine group, as compared with the saline group (Fig. 2G; *t*_(18)_ = 2.97, *p* < 0.001). This result further indicates that repeated morphine administration causes desensitization of MOR-mediated inhibition of synaptic transmission in the dMSN→GPh pathway.

### Noncontingent opioid recruits dMSN ensembles and contingent opioid potentiates striatopallidal GABAergic transmission in GPh neurons

Having shown that repeated MOR activation by non-contingent morphine injection enhances GABAergic dMSN→GPh transmission, we next examined whether this transmission was altered by chronic exposure to a more potent and prevalent MOR agonist, fentanyl. We first tested whether repeated fentanyl injections also induced the hyperlocomotion observed using morphine. C57BL/6 mice received daily fentanyl injections for 5 d (on days 3–7) and then on days 16 and 17. Locomotor activity was measured immediately after each injection. The injections on days 16 and 17 were administered to test whether fentanyl withdrawal caused locomotor sensitization (<u>Grimm et al., 2001</u>; <u>Li et al., 2008</u>; <u>Pickens et al., 2011</u>; <u>Reiner et al., 2019</u>). We found that fentanyl induced hyperactivity after each injection (Fig. 3A, B; *F*_(1,8)_ = 19.84, *p* < 0.01) and that there was no sensitization during the first 5-d series of injections (Fig. 3A; D3 versus D4–D7: *q* = 0.75–2.21, *p* > 0.05), which was consistent with the morphine effects (Fig. 2B). However, locomotor activity was significantly higher on day 16 than on day 7 (Fig. 3C; *t*_(5)_ = -2.66, *p* = 0.045). The fentanyl injections administered after the 8-d withdrawal period produced a longer period of hyperlocomotion (1 h versus 40 min) (Fig. 3D). These results suggest that non-contingent fentanyl administration also increases locomotor activity and that fentanyl withdrawal causes locomotor sensitization.

**Figure 3.**
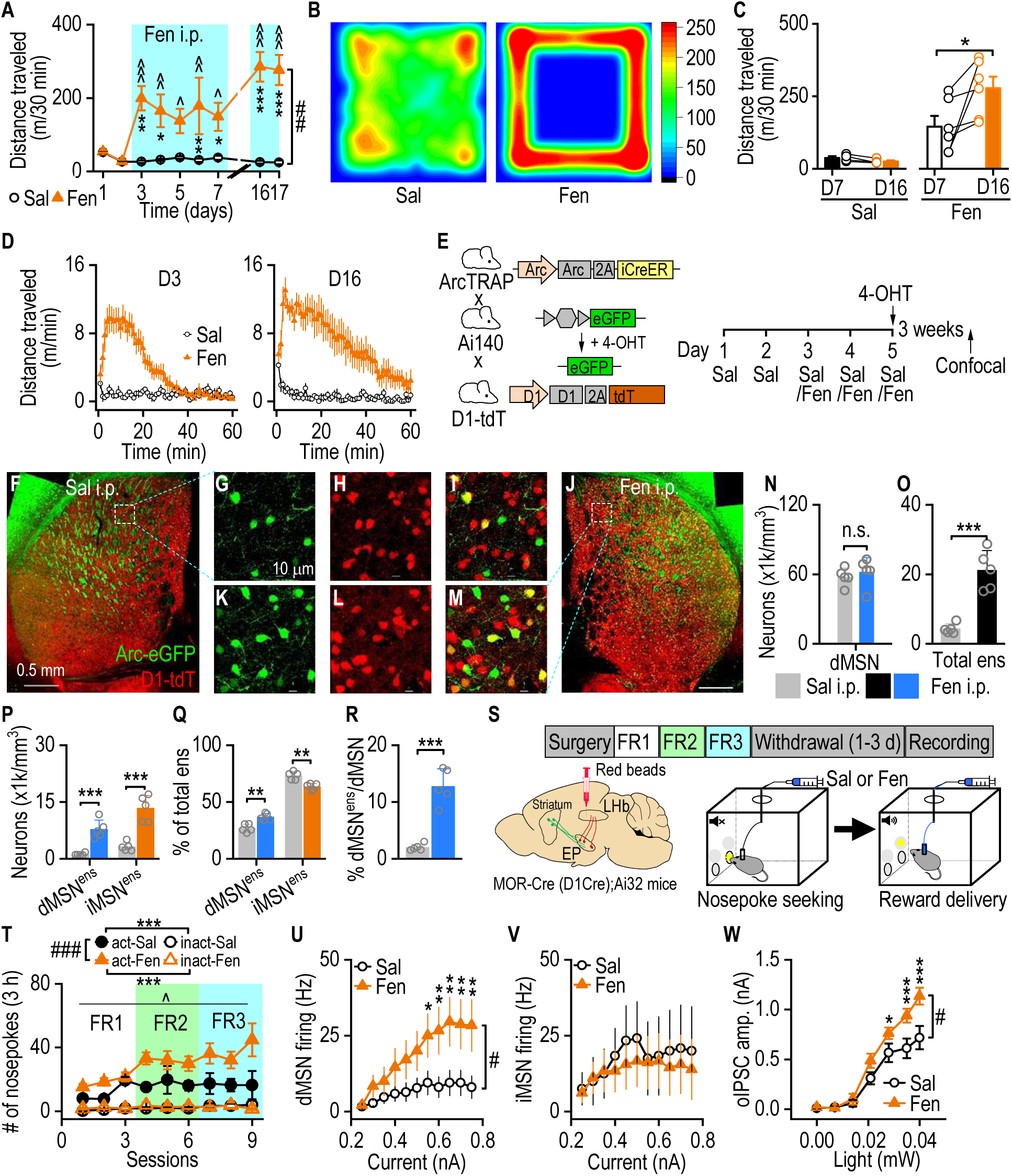
Intravenous fentanyl administration potentiates striatopallidal GABAergic transmission in GPh neurons. ***A***, Time-course of i.p. fentanyl (Fen, 0.3 mg/kg)-induced hyperactivity in C57BL/6 mice; ^##^*p* < 0.01; \**p* < 0.05, \*\**p* < 0.01, \*\*\**p* < 0.001 versus saline (Sal) at the same days; ^^*p* < 0.01, ^^^*p* < 0.001 versus days 1 and 2. ***B***, Representative heatmap showing the locomotor activity following i.p. fentanyl versus saline on day 3. ***C***, Comparison of locomotor activity on day 16 versus day 7; \**p* < 0.05. ***D***, Representative time-courses of fentanyl-induced hyper-locomotor activity on days 3 (left) and 16 (right). Note the slower decay on day 16 than on day 3. ***E***, Schematics showing the experimental design used to Arc-TRAP fentanyl-induced dMSN ensembles (dMSN^ens^). Left: ArcTRAP;Ai140;D1-tdT mice expressed tdT in dMSNs and injection of 4-hydroxytamoxifen (4- OHT) induced GFP expression in TRAPed neurons. Right: Timeline for the experiment. ***F*–*M***, Sample images of ensembles recruited following i.p. administration of saline (*F–I*) or fentanyl (*J*– *M*) (green) and dMSNs (red) in the DMS. dMSN^ens^ expressed GFP and tdT (yellow; *I*, *M*). ***N, O***, Although i.p. fentanyl did not alter the dMSN density (*O*), it did increase the total ensemble density (ens; *P*); n.s. not significant, *p* > 0.05; \*\*\**p* < 0.001. ***P***, Fentanyl administration increased the densities of both dMSN^ens^ and putative iMSN^ens^ (tdT-negative ensembles); \*\*\**p* < 0.001. ***Q***, Expressed as a percentage of total ensembles, the proportion of dMSN^ens^ increased, while the proportion of iMSN^ens^ decreased following i.p. fentanyl; \*\**p* < 0.01. ***R***, Fentanyl administration increased the percentage of total dMSNs recruited into ensembles; \*\*\**p* < 0.001. ***S***, Schematic showing the infusion of red beads, intravenous self-administration (IVSA) of fentanyl, and the experimental timeline. The LHb of MOR-Cre (or D1-Cre);Ai32 mice was infused with red beads prior to training the animals to self-administer fentanyl (Fen, 3 μg/kg) or saline (Sal, 0.9%) using FR1–3 schedules. Slice recordings were conducted after 1–3 d of withdrawal from the last IVSA session. ***T***, Significantly more active nosepokes (act) were observed in the fentanyl group than in the saline group; ^###^*p* < 0.001 for fentanyl versus saline (active nosepokes); ^^^*p* < 0.05 for different sessions; \*\*\**p* < 0.001 for active versus inactive (inact) nosepokes within each treatment. ***U*, *V***, Fentanyl IVSA increased the frequency of evoked firing in dMSNs (*U*) but not in iMSNs (*V*) from D1-Cre;Ai32 mice; ^#^*p* < 0.05; \**p* < 0.05, \*\**p* < 0.01 versus the saline group at the same stimulating intensity. ***W***, Fentanyl IVSA increased oIPSC amplitudes in GPh neurons from MOR-Cre;Ai32 and D1-Cre;Ai32 mice; ^#^*p* < 0.05; \**p* < 0.05, \*\*\**p* < 0.001 versus the saline group at the same stimulating intensity. Two-way RM ANOVA followed by Tukey *post hoc* test for *A* and *U*–*W*, paired *t* test for *C*, unpaired *t* test for *N–R*, and three-way ANOVA for *T*; n = 5/2 mice per group (*N–R*), 12/3 (*U*, Sal), 8/3 (*U*, Fen), 4/3 (*V*, Sal), 6/3 (*V*, Fen), 10/5 (*W*, Sal), and 18/8 (*W*, Fen).

Striatal dMSN excitation is known to increase motor functions (<u>Kravitz et al., 2010</u>). Thus, we examined whether non-contingent fentanyl exposure recruited dMSN ensembles (dMSN^ens^). To achieve this goal, we generated ArcTRAP;Ai140;D1-tdT mice in which dMSNs expressed tdT and TRAPed neuronal ensembles expressed GFP in response to 4-hydroxytamoxifen (4-OHT) injection (<u>DeNardo et al., 2019</u>) (Fig. 3E). Three weeks after TRAPing, we examined fentanyl-recruited GFP-positive ensembles and their overlap with tdT-positive dMSNs or putative tdT- negative iMSNs (Fig. 3F–M). While not altering the dMSN density in the DMS as expected (Fig. 3N; *t*_(8)_ = -0.63, *p* > 0.05), fentanyl injection activated significantly more ensembles, as compared to saline administration (Fig. 3O; *t*_(8)_ = -6.40, *p* < 0.001). Interestingly, although both dMSN^ens^ and iMSN ensemble (iMSN^ens^) densities were increased in the fentanyl mice (Fig. 3P; dMSN^ens^: *t*_(8)_ = -6.05, *p* < 0.001; iMSN^ens^: *t*_(8)_ = -6.28, *p* < 0.001), the proportion of total ensembles that were dMSN^ens^ was significantly greater in the fentanyl group than in the saline group (Fig. 3Q; dMSN^ens^: *t*_(8)_ = -4.72, *p* < 0.01; iMSN^ens^: *t*_(8)_ = 4.72, *p* < 0.01). Importantly, we found that fentanyl exposure significantly increased the percentage of dMSN^ens^ (out of total dMSNs), as compared to saline injections (Fig. 3R; *t*_(8)_ = -7.57, *p* < 0.001). These results indicate that non-contingent fentanyl administrations recruit dMSN^ens^ in the DMS, an effect that may account for fentanyl-induced hyperlocomotion.

Next, we examined whether contingent fentanyl exposure altered striatopallidal activity as morphine (Fig. 2). To achieve this, we first trained C57BL/6 mice for IVSA of fentanyl in operant chambers. We found that fentanyl reinforced operant self-administration (Supplementary Fig. 2A– C; A: *F*_(11,11)_ = 4.36, *p* < 0.05; *q* = 4.93, *p* < 0.05 (FR3_3); *q* = 5.56–8.07, *p* < 0.01; B: *F*_(11,21)_ =0.83, *p* > 0.05; C: *F*_(11,21)_ = 1.25, *p* > 0.05). The dose-response curve exhibited an “inverted U” shape, with the highest active responses at 0.03 µg/kg (Supplementary Fig. 2D; nosepokes: *F*_(3,13)_ = 3.15, *p* = 0.06; infusion: *F*_(3,13)_ = 2.34, *p* = 0.12). We therefore used this dose for all subsequent IVSA experiments. We next trained MOR-Cre (or D1-Cre);Ai32 mice for IVSA of fentanyl or saline (Fig. 3S). The LHb of these mice were pre-infused with red beads to facilitate selective recording of GPh neurons (Fig. 3S). Mice were able to self-administer either fentanyl or saline (<u>Li et al., 2015</u>) (Fig. 3T; *F*_(1,162)_ = 185.67, *p* < 0.001). However, active nosepokes for fentanyl were significantly higher than those for saline (Fig. 3T; *F*_(1,162)_ = 10.23, *p* < 0.01). Additionally, the accuracy of both the fentanyl and saline groups was higher than 75% (Supplementary Fig. 2E; *F*_(1,9)_ = 0.998, *p* = 0.34). Significantly more fentanyl infusions were administered (> 10 across the training schedules) than saline infusions (Supplementary Fig. 2F; *F*_(1,9)_ = 5.22, *p* < 0.05). One to three days after the last IVSA session, we prepared slices to measure dMSN activity and oIPSCs in GPh neurons. We observed that contingent fentanyl exposure significantly increased the excitability of dMSNs, but not of iMSNs, in the DMS (dMSNs: Fig. 3U, *F*_(1,18)_ = 5.27, *p* < 0.05; iMSNs: Fig. 3V, *F*_(1,8)_ = 0.11, *p* > 0.05). In GPh slices, oIPSC amplitudes were significantly greater in the fentanyl group than in the saline group (Fig. 3W; *F*_(1,26)_ = 6.93, *p* < 0.05), which was consistent with the morphine injection results (Fig. 2D). Taken together, these results demonstrate that contingent fentanyl administrations enhance dMSN activity and potentiate striatopallidal GABAergic transmission in GPh neurons.

### Contingent opioid potentiates GABAergic transmission in DMS patch neurons but not in matrix neurons

In addition to sending their MOR-expressing axons to the GPh, dMSNs send collaterals to innervate other striatal neurons. MORs were mainly expressed in the patch (but not matrix) compartment in double transgenic (MOR-Cre;Snap25-GFP) mice (Fig. 4A). Notably, using a triple transgenic (MOR-Cre;Snap25-GFP;D1-tdT) line with MOR^+^ neurons expressing GFP and dMSNs containing tdT, we found that MOR^+^ dMSNs were predominantly expressed in the patch compartment (Fig. 4B; patches: Fig. 4C-left, *F*_(2,66)_ = 254.39, *p* < 0.001; Supplementary Fig. 3A, *F*_(2,66)_ = 811.22, *p* < 0.001), but not in the matrix compartment (Fig. 4B; matrix: Fig. 4C-right, *F*_(2,66)_ = 300.47, *p* < 0.001; Supplementary Fig. 3B, *F*_(2,63)_ = 2940.2, *p* < 0.001). These MOR^+^ dMSNs projected to the EP and SNr (<u>Stephenson-Jones et al., 2016</u>) (Supplementary Fig. 3C, D). We confirmed that DAMGO suppressed electrically evoked IPSCs in patch neurons, but not in matrix neurons (<u>Banghart et al., 2015b</u>; <u>Miura et al., 2007</u>) (Supplementary Fig. 3E; Supplementary Fig. 3F: *F*_(1,10)_ = 198.63, *p* < 0.001; Supplementary Fig. 3G: *t*_(5)_ = 6.25, *p* < 0.01). Then, we examined whether fentanyl IVSA altered GABAergic transmission in patch and matrix neurons using a transgenic line (MOR-Cre;Ai32) in which MOR^+^ neurons expressed ChR2-EYFP (Fig. 4D).

**Figure 4.**
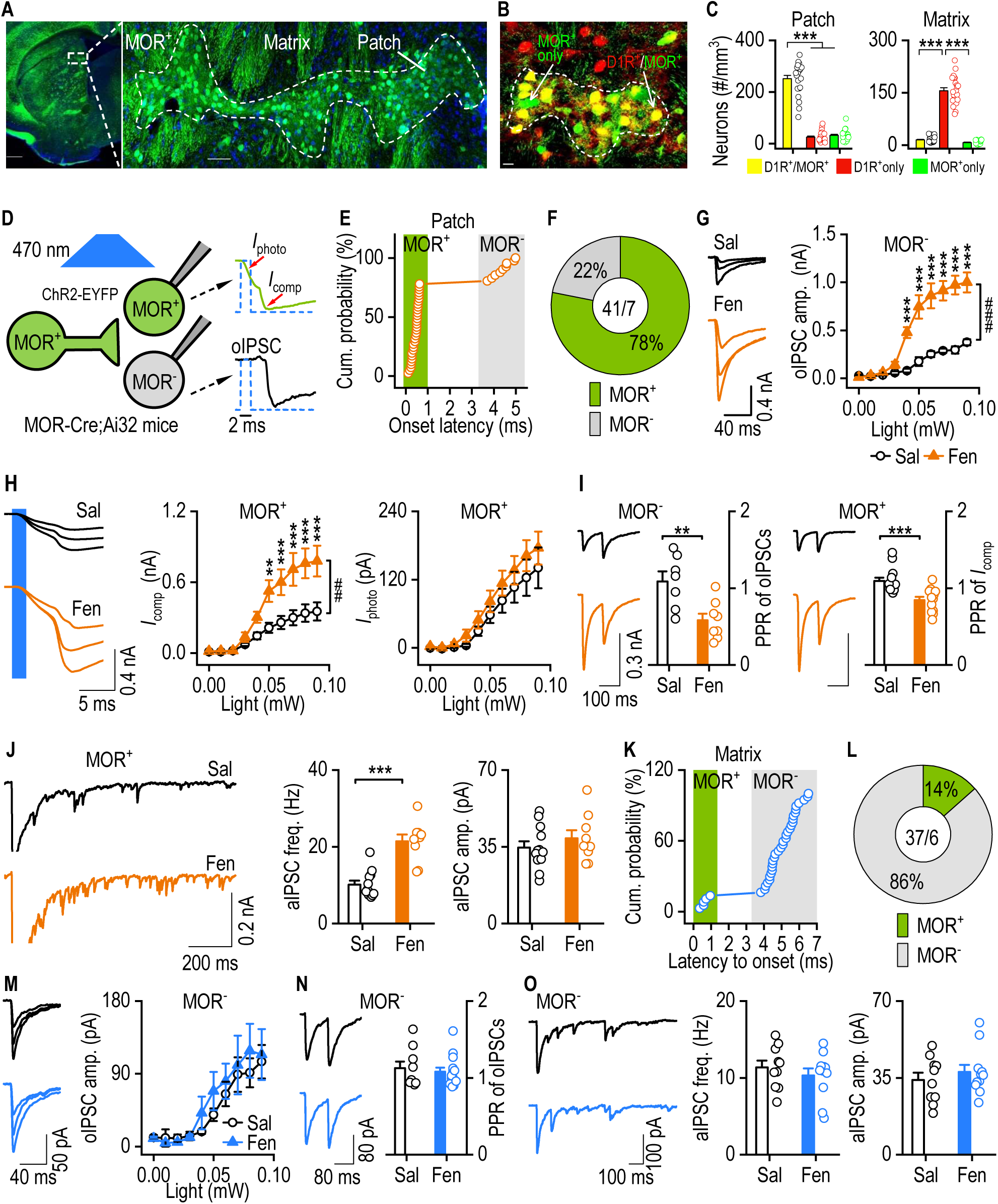
Fentanyl IVSA potentiates GABAergic transmission in patch, but not matrix, neurons in the DMS. ***A***, Representative images showing the structural separation of patch and matrix compartments in MOR-Cre;Snap25-GFP;D1-tdT mice. Scale bar, 1 mm (left), 0.1 mm (right). ***B***, Representative image showing the colocalization (yellow) of MOR (green) and D1R (red) expression in the patch and matrix compartments. Scale bar, 0.1 µm. ***C***, The patch, but not matrix, compartment contained mainly MOR-expressing dMSNs (GFP^+^ and tdT^+^); \*\*\**p* < 0.001. ***D***, Diagram with examples of recordings in MOR^+^ and MOR^-^ neurons in DMS slices. Optogenetic excitation of MOR^+^ neurons induced two types of responses with distinct onset latencies: oIPSC only (latency > 2 ms; MOR^-^ neurons) and direct depolarization (latency < 2 ms, *I*_photo_) plus subsequent synaptic transmission (compound currents, *I*_comp_; MOR^+^ neurons). ***E***, Separation of the onset latencies of responses from MOR^+^ and MOR^-^ neurons in the patches. ***F***, The majority of patch neurons were MOR^+^. ***G***, Fentanyl IVSA significantly increased the oIPSC amplitudes in MOR^-^ neurons within patches; ^###^*p* < 0.001; \*\*\**p* < 0.001 versus the saline group at the same stimulating intensity. ***H***, Fentanyl IVSA significantly increased *I*_comp_ (left and middle) without changing *I*_photo_ (left and right) in patch MOR^+^ neurons; ^##^*p* < 0.01; \*\**p* < 0.01 and \*\*\**p* < 0.001 versus the saline group at the same stimulating intensity. ***I***, Fentanyl IVSA significantly decreased PPRs in patch MOR^-^ (left) and MOR^+^ (right) neurons; \*\**p* < 0.01, \*\*\**p* < 0.001. ***J***, Fentanyl IVSA increased the frequency, but not the amplitude, of asynchronous IPSCs (aIPSCs; recorded) in MOR^+^ neurons; \*\*\**p* < 0.001. ***K***, Separation of response onset latencies in matrix neurons that were MOR^+^ or MOR^-^. ***L***, The majority of matrix neurons were MOR^-^. ***M–O***, Fentanyl IVSA did not alter the oIPSC amplitude (*M*), PPR (*N*), or the aIPSC frequency or amplitude (*O*) in MOR^-^ matrix neurons. MOR-Cre;Ai32 mice were used in *D–O*. Two-way RM ANOVA followed by Tukey *post hoc* test (*G*, *H*, *M*), one way ANOVA (*C*), and unpaired *t* test (*I*, *J*, *N*, *O*); n = 23/2 (*C,* left), 23/2 (*C,* right), 41/7 (*E*, *F*), 8/4 (*G*, Sal), 10/3 (*G*, Fen), 12/3 (*H*-middle, Sal), 10/3 (*H*-middle, Fen), 12/3 (*H*-right, Sal), 14/3 (*H*- right, Fen), 7/3 (*I-left*, Sal), 8/3 (*I-left*, Fen), 14/4 (*I*-right, Sal), 13/3 (*I*-right, Fen), 12/3 (*J*, Sal), 9/2 (*J*, Fen), 37/6 (*K*, *L*), 13/4 (*M*, Sal), 12/3 (*M*, Fen), 9/4 (*N*, Sal), 14/2 (*N*, Fen), 10/3 (*O*, Sal), and 11/3 (*O*, Fen).

Optically induced responses were recorded after 1–3-d withdrawal from fentanyl IVSA (Fig. 3S). Blue light stimulation induced two types of optical responses with different latencies: < 2 ms or > 2 ms (Fig. 4D, E). The response with a latency of < 2 ms was likely mediated by ChR2-induced direct depolarization (*I*_photo_) of MOR^+^ neurons. In contrast, those with a latency of > 2 ms reflected synaptic transmission (oIPSCs) in MOR-negative (MOR^-^) neurons. In the patches, we found that ∼78% of recorded neurons were MOR^+^ and ∼22% were MOR^-^ (Fig. 4F); this is consistent with a previous report (<u>Smith et al., 2016b</u>) and our histological findings (Fig. 4C). In MOR^-^ neurons, fentanyl IVSA significantly increased the oIPSC amplitudes (Fig. 4G; *F*_(1,16)_ = 33.79, *p* < 0.001). Similarly, in MOR^+^ neurons, fentanyl IVSA enhanced the peak amplitude of optically induced compound responses (Fig. 4H left and middle; *F*_(1,20)_ = 10.00, *p* < 0.01), which comprised ChR2- mediated direct depolarization and synaptic responses. However, fentanyl IVSA did not alter direct depolarization-induced responses measured at 2, 3, or 4 ms from the start of the light stimulation (Fig. 4H right; *F*_(1,24)_ = 1, *p* > 0.05; Supplementary Fig. 3H), suggesting that fentanyl IVSA enhanced synaptic transmission in MOR^+^ neurons. PPRs in both MOR^+^ and MOR^-^ neurons were significantly higher in the fentanyl group than in the saline controls (MOR^+^: Fig. 4I left, *t*_(13)_ = 3.28, *p* < 0.01; MOR^-^: Fig. 4I right, *t*_(25)_ = 4.23, *p* < 0.001). We consistently found that the frequency, but not the amplitude, of asynchronized inhibitory postsynaptic currents (aIPSCs) in MOR^+^ neurons was significantly higher in fentanyl-treated mice than in the saline controls (Fig. 4J; frequency: *t*_(19)_ = 5.9, *p* < 0.001; amplitude: *t*_(19)_ = 0.94, *p* > 0.05). These results suggest that contingent fentanyl IVSA potentiates MOR-expressing outputs onto patch neurons, at least in part via a presynaptic mechanism.

We also observed two types of light-induced responses in matrix neurons; similar to those found in patches (Fig. 4E), these had different latencies (Fig. 4K). In contrast to the patch neurons (Fig. 4E, F), we found that only 14% of recorded matrix neurons were MOR^+^ (with a latency of < 2 ms) and 86% were MOR^-^ (with a latency of > 2 ms) (Fig. 4L). Interestingly, fentanyl IVSA did not alter the oIPSC amplitudes or PPRs in matrix MOR^-^ neurons (oIPSC amplitude: Fig. 4M, *F*_(1,23)_ = 0.36, *p* > 0.05; PPR: Fig. 4N, *t*_(21)_ = 0.47, *p* > 0.05). The aIPSC frequencies and amplitudes in matrix MOR^-^ neurons were also unaffected by fentanyl IVSA (Fig. 4O; frequencies: *t*_(19)_ = 0.82, *p* > 0.05; amplitudes: *t*_(19)_ = 0.8, *p* > 0.05). Overall, these results indicate that fentanyl IVSA potentiates GABAergic transmission in the patch, but not matrix, compartments.

### Contingent opioid potentiates striatonigral GABAergic transmission in SNc dopaminergic neurons

In addition to their connections with the striatum and EP, MOR-expressing striatal neurons send axons to innervate midbrain dopaminergic neurons (<u>Crittenden et al., 2016</u>; <u>Evans et al., 2020</u>; <u>Fujiyama et al., 2011</u>; <u>Watabe-Uchida et al., 2012</u>). To investigate this innervation, we infused AAV1- Cre into the DMS of Ai14 mice. AAV1-Cre is known to induce anterograde transsynaptic expression of Cre (<u>Chan et al., 2017</u>; <u>Yao et al., 2018</u>; <u>Zhang et al., 2018</u>; <u>Zingg et al., 2017</u>; <u>Zingg et al.,</u> <u>2020</u>). We observed robust levels of tdT in the SNc (Fig. 5A). Since dMSNs project to the SNc and iMSNs do not (<u>Cheng et al., 2017b</u>; <u>Crittenden et al., 2016</u>; <u>Fujiyama et al., 2011</u>; <u>Gerfen and</u> <u>Surmeier, 2011</u>; <u>Kreitzer and Malenka, 2008</u>; <u>Xiao et al., 2020</u>), this finding suggests that dMSNs project to SNc neurons. We then tested whether fentanyl IVSA altered dMSN-derived GABAergic inputs onto SNc dopaminergic neurons in D1-Cre;Ai32 mice, in which dMSNs expressed ChR2-EYFP (<u>Lu et al.,</u> <u>2019</u>) (Fig. 5B). In this experiment, we selected D1-Cre rather than MOR-Cre mice to avoid activating other MOR-expressing GABAergic inputs (<u>Galaj et al., 2020</u>; <u>Matsui et al., 2014</u>). SNc dopaminergic neurons were identified by the presence of hyperpolarization-induced cation currents (*I*_h_), and GABA_A_R-IPSCs were confirmed using picrotoxin (Fig. 5C). We found that fentanyl IVSA significantly potentiated GABAergic transmission in SNc dopaminergic neurons after 1–3-d withdrawal following the last IVSA (Fig. 5D, *F*_(1,35)_ = 16.82 *p* < 0.001). Moreover, these increased inhibitory currents were accompanied by significantly lower PPRs (Fig. 5E, *t*_(33)_ = 5.05, *p* < 0.001) and elevated aIPSC frequencies, while amplitudes were unaffected (Fig. 5F; frequencies: *t*_(34)_ = -7.02, *p* < 0.001; amplitudes: *t*_(34)_ = -0.43, *p* > 0.05). These results indicate that contingent fentanyl IVSA enhances D1R-expressing inputs onto SNc dopaminergic neurons via a presynaptic mechanism.

**Figure 5.**
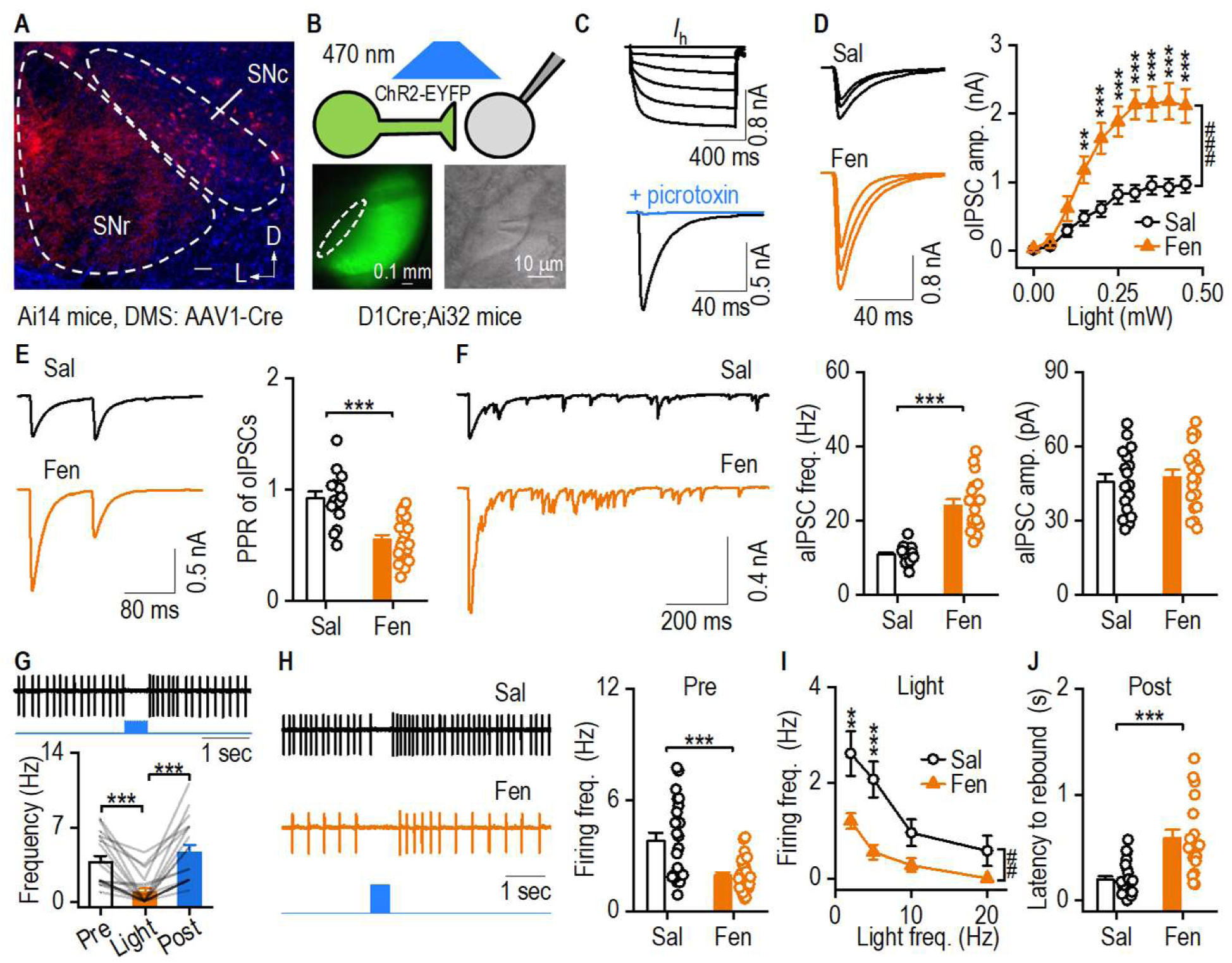
Fentanyl IVSA potentiates GABAergic transmission from striatal dMSNs to SNc dopaminergic neurons. D1-Cre;Ai32 mice were trained with fentanyl or saline IVSA, and spontaneous firing and oIPSCs were recorded in SNc dopaminergic neurons 1–3 d after the last fentanyl administration. ***A***, Confocal image of the midbrain showing that SNc neurons received inputs from DMS dMSNs. AAV1-Cre was infused into the DMS of Ai14 mice. Scale bar: 0.1 mm. D, dorsal; L, lateral. ***B***, Schematic (top) and sample images (bottom) showing stimulating and recording in the SNc of D1-Cre;Ai32 mice. **C**, Sample traces of *I*_h_ currents (top) and oIPSCs (bottom) in SNc dopaminergic neurons. ***D***, Fentanyl IVSA significantly enhanced oIPSC amplitudes; ^###^*p* < 0.001; \*\**p* < 0.01, \*\*\**p* < 0.001 versus the saline group at the same stimulating intensities. ***E***, Fentanyl IVSA inhibited oIPSC PPRs in SNc dopaminergic neurons; \*\*\**p* < 0.001. ***F***, Fentanyl IVSA potentiated aIPSC frequencies (left and middle), but not amplitudes (left and right); \*\*\**p* < 0.001. ***G***, Optogenetic stimulation of D1R^+^ inputs (10 pulses, 20 Hz) reversibly suppressed spontaneous firing of SNc dopaminergic neurons; \*\**p* < 0.01, \*\*\**p* < 0.001. ***H***, Fentanyl IVSA reduced baseline spontaneous firing in SNc dopaminergic neurons; \*\*\**p* < 0.001. ***I***, Fentanyl IVSA reduced D1R^+^ input-mediated suppression of spontaneous firing in SNc dopaminergic neurons; ^##^*p* < 0.01; \**p* < 0.05, \*\**p* < 0.01, \*\*\**p* < 0.001 versus the saline group at the same stimulating intensities. ***J***, Fentanyl IVSA enhanced the latency of rebound firing after light stimulation; \*\*\**p* < 0.001. One-way (*G*) or two-way (*J*, *D*) RM ANOVA followed by Tukey *post hoc* test; unpaired *t* test (*E*, *F*, *H*, *J*). n = 18/3 (*D;* Sal), 19/5 (D; Fen), 15/3 (*E*; Sal), 20/5 (*E*; Fen), 17/3 (*F*; Sal), 19/5 (*F*; Fen), 18/3 (*G*), 23/4 (*H*; Sal), 27/4 (*H*; Fen), 14/2 (*I*; Sal), 11/3 (*I*; Fen), 23/3 (*J*; Sal), 19/5 (*J*; Fen).

Since GABAergic transmission is inhibitory, we tested whether optogenetic stimulation of D1R-expressing fibers reduced the spontaneous firing of SNc dopaminergic neurons. We discovered that light stimulation reversibly suppressed the firing frequencies of these neurons (Fig. 5G; *F*_(2,34)_ = 34.147, *p* < 0.001; Supplementary Fig. 4A: *F*_(2,34)_ = 142.01, *p* < 0.001). Interestingly, baseline firing rates were significantly lower in fentanyl-treated mice than in saline controls (Fig. 5H, *t*_(48)_ = 4.39, *p* < 0.001). Notably, fentanyl IVSA significantly reduced D1R^+^ input- mediated suppression of spontaneous firing, as compared to saline IVSA (Fig. 5I; *F*_(1.23)_ = 10.16, *p* < 0.01; Supplementary Fig. 4B: *F*_(1,23)_ = 15.2, *p* < 0.001). In addition, fentanyl IVSA markedly increased the latency of rebound firing following light stimulation (Fig. 5J, *t*_(40)_ = -4.91, *p* < 0.001). However, the intrinsic activity of SNc dopaminergic neurons was not altered by fentanyl exposure (Supplementary Fig. 4C, evoked firing: *F*_(1,25)_ = 0.79, *p* > 0.05; Fig. 4D, sag amplitudes: *F*_(1,24)_ = 2.28, *p* > 0.05). Taken together, our results suggest that contingent fentanyl IVSA enhances GABAergic outputs from dMSNs onto SNc dopaminergic neurons; this strengthens their suppression of spontaneous firing and thus reduces the baseline firing rate of these neurons.

### Opioid withdrawal induces depression-like behaviors and reinstatement

Having shown that chronic fentanyl exposure reduces the spontaneous firing of dopaminergic neurons, an effect that is associated with negative emotional states (<u>Koob, 2021</u>), we next examined whether withdrawal from repeated fentanyl injections induced depression-like behaviors. C57BL/6 mice received daily fentanyl (0.3 mg/kg, i.p.) injections for 2 weeks, and behavioral tests were conducted on withdrawal day 10. We found that fentanyl withdrawal increased immobility time (Fig. 6A; *t*_(26)_ = -2.47, *p* < 0.05) and decreased distance traveled (Fig. 6B: *t*_(26)_ = -2.74, *p* < 0.05; Fig. 6C: *F*_(1,26)_ = 6.31, *p* < 0.05) in the open field test. These behavioral differences may reflect increased levels of depression following fentanyl withdrawal. To explore this possibility, we conducted sucrose splash tests. Fentanyl withdrawal caused a trend towards increased latency until the first grooming (Fig. 6D; *t*_(25)_ = -1.99, *p* = 0.0583) and significantly decreased grooming frequency (Fig. 6E; *t*_(25)_ = -2.08, *p* < 0.05) and duration (Fig. 6F, *t*_(25)_ = 2.27, *p* < 0.05). These results suggest that withdrawal from non-contingent fentanyl administration induces depression-like behaviors. Unbiased cluster analysis (<u>Li et al., 2021b</u>) of the open field tests, social interaction tests (Supplementary Fig. 5), and sucrose splash tests yielded two clusters: non-depression-like and depression-like behaviors (Fig. 6G and H). We found that fentanyl withdrawal marginally increased the percentage of animals showing depression-like behaviors (Fig. 6I, *p* = 0.054). Withdrawal-induced negative emotional states such as depression promote relapse of drug use (<u>Koob, 2021</u>). Next, we used conditioned place preference (CPP) to examine whether long-term withdrawal from fentanyl injections elicited reinstatement (<u>Cunningham</u> <u>et al., 2006</u>; <u>Milton and Everitt, 2012</u>). Animals received fentanyl injections once daily for 7 consecutive days, and CPP testing was conducted on days 8 and 36 (Fig. 6J). We observed that fentanyl caused a robust CPP on day 8 (Fig. 6K; WD1: *t*_(16)_ = -3.48, *p* < 0.01). After 28 d of abstinence, fentanyl-treated animals exhibited a strong context-induced preference for the drug- paired chamber, comparable to the level at 1-d withdrawal (Fig. 6K; WD28: *t*_(16)_ = -3.10, *p* < 0.01). These data suggest that negative emotions induced by withdrawal from non-contingent fentanyl exposure drive reinstatement. Lastly, we tested whether animals that received contingent fentanyl administrations also exhibited reinstatement (Fig. 6L). We found that extinction training significantly inhibited fentanyl seeking (Fig. 6L; *F*_(5,10)_ = 9.61, *p* < 0.01; *q* = 5.32, *p* < 0.05; *q* = 7.47–11.5, *p* < 0.001). After extinction, however, fentanyl-paired cues strongly promoted the reinstatement of fentanyl-seeking behaviors (Fig. 6L; Reinst vs. last Ext: *q* = 8.19, *p* < 0.001). Taken together, our data indicate that withdrawal from opioid exposure induced negative emotional states, driving the reinstatement of drug-seeking behaviors.

**Figure 6.**
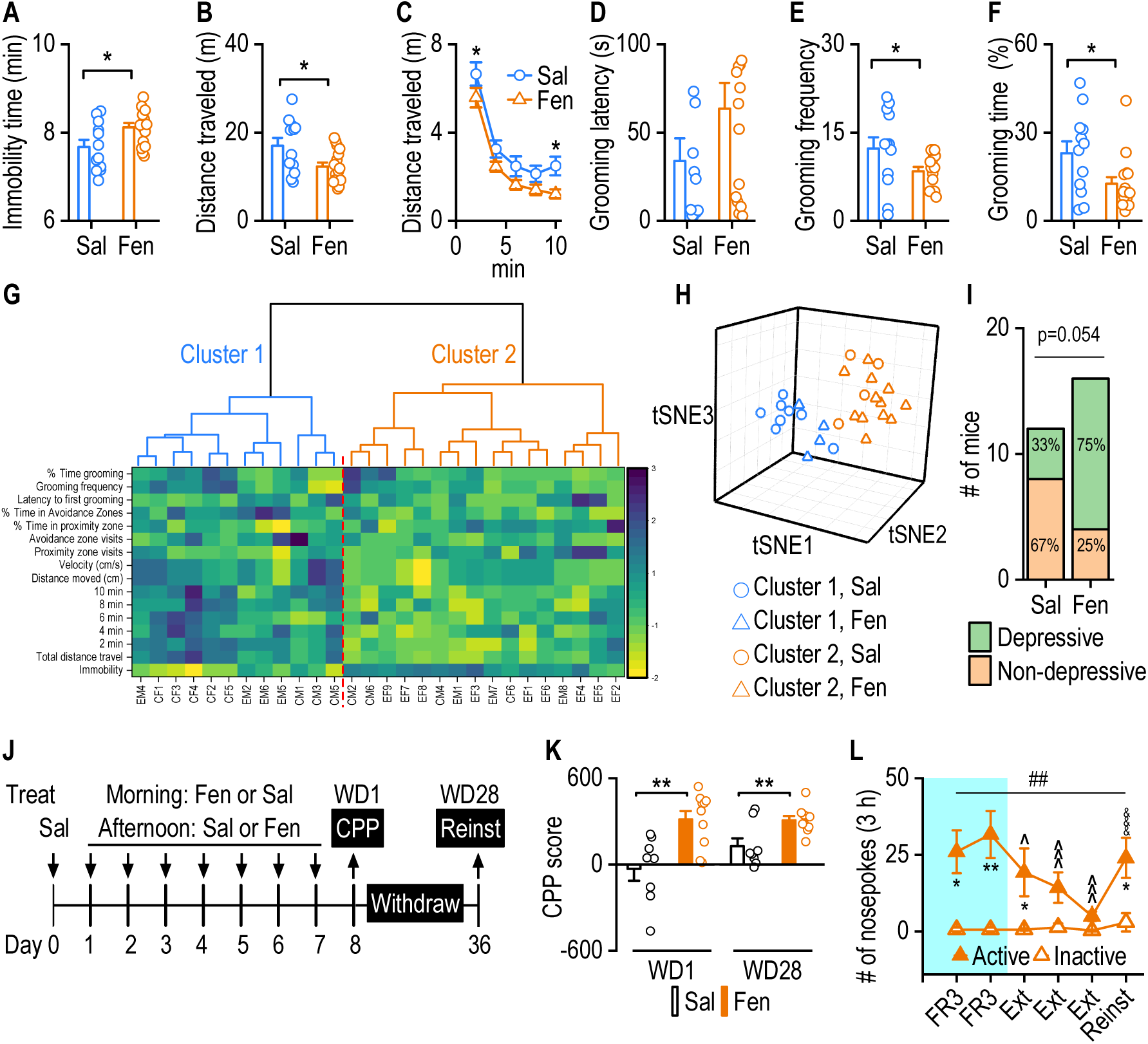
Repeated fentanyl administration induces depression-like behaviors and relapse. C57BL/6 mice received daily i.p. fentanyl injections for 14 d, and behavioral tests were conducted 10 d after the last injection. ***A–C***, Fentanyl withdrawal significantly increased immobility time (*A*) and decreased the distance traveled in a 10-min open field test (*B*, *C*); \**p* < 0.05. ***D–F***, In the sucrose splash test, repeated i.p. fentanyl did not alter the latency to the first grooming (*D*) but did reduce grooming frequency (*E*) and the percentage of grooming time (*F*); \**p* < 0.05. ***G***, Fentanyl withdrawal induced two behavioral phenotypes, identified using hierarchical cluster analysis with t-distributed stochastic neighbor embedding (tSNE) projection of data from multiple behavioral tests. ***H***, Three-dimensional tSNE representation of clusters of non-depression-like (blue; cluster 1) and depression-like (orange; cluster 2) behaviors. ***I***, Fentanyl withdrawal induced a trend toward an increased percentage of animals displaying depression-like behaviors; *p* = 0.053, Fisher’s exact test. ***J***, Experimental timeline of fentanyl (0.03 mg/kg)-induced conditioned place preference (CPP) and context-induced reinstatement (Reinst) in Long-Evans rats. Saline (1 ml/kg); WD1 and WD28, withdrawal day 1 and 28, respectively. ***K***, Repeated fentanyl injections induced a robust CPP and context-induced reinstatement of CPP. The CPP score was calculated by subtracting the time spent in the non-paired chamber from the time spent in the fentanyl-paired chamber; \*\**p* < 0.01. ***L***, Fentanyl IVSA caused cue-induced reinstatement of drug-seeking. ^##^*p* < 0.01 for sessions; \**p* < 0.05, \*\**p* < 0.01 versus inactive nosepokes during the same session; ^*p* < 0.05, ^^*p* < 0.01, ^^^*p* < 0.001 versus FR3 for active nosepokes; ^&&&^*p* < 0.001 versus the last extinction (Ext) for active nosepokes by Tukey *post hoc* test. Unpaired *t* test (*A, B, D–F, I, K*), and two-way RM ANOVA (*C*, *L*); n = 12 (*A–I*; Sal), 16 (*A–I;* Fen), 8 (*K*; Sal), 10 (*K*; Fen), 3 (*L*).

## DISCUSSION

This study demonstrated that although acute MOR activation suppressed GABAergic striatopallidal (dMSN→GPh) transmission, repeated exposure to morphine or fentanyl potentiated this transmission. In addition, fentanyl IVSA potentiated GABAergic transmission within the striatal patch, but not the matrix, compartment. Fentanyl IVSA also potentiated GABAergic striatonigral (dMSN→SNc) transmission and suppressed SNc dopaminergic firing. Importantly, fentanyl withdrawal induced depression-like behaviors and caused reinstatement. These data suggest that repeated opioid use potentiates GABAergic striatopallidal and striatonigral transmission; the resulting reduction in dopaminergic activity may contribute to the negative emotional states that trigger relapse.

### Acute MOR activation suppresses GABAergic striatopallidal transmission

The addictive properties of opioids are mediated by MORs, and MOR activation is known to inhibit neuronal activity in the hippocampus and ventral tegmental area (<u>Al-Hasani and Bruchas,</u> <u>2011</u>; <u>Evans, 2004</u>; <u>Gaveriaux-Ruff and Kieffer, 2002</u>; <u>Gopalakrishnan et al., 2021</u>; <u>Volkow et al., 2019</u>; <u>Williams et al., 2001</u>; <u>Williams et al., 2013</u>). Our study found that acute MOR activation inhibited GABAergic dMSN→EP/GPh transmission. EP neurons receive GABAergic inputs from striatal dMSNs and the GPe (<u>Bolam and Smith, 1992</u>; <u>Deniau et al., 1978</u>). Since D1 receptors are only present in dMSN inputs (<u>Lavian et al., 2018</u>), Chrimson-mediated oIPSCs were mainly from dMSN terminals. Furthermore, our data also suggested that most of these dMSN afferents originated from the patch compartment because only the patch or exo-patch compartment neurons expressed MORs (<u>Brimblecombe and Cragg, 2017</u>; <u>Gerfen, 1984</u>; <u>Smith et al., 2016a</u>). Within the EP, GPh neurons are located in a complex pathway that connects striatal dMSNs with the LHb, rostromedial tegmental nucleus (RMTg), and SNc dopaminergic neurons (<u>Baker et al., 2016</u>; <u>Mollick et al., 2020</u>). During negative emotional states, GPh projections excite the LHb, which regulates negative valence and drives aversive behaviors (<u>Faget et al., 2018</u>; <u>Li et al., 2021a</u>; <u>Meye</u> <u>et al., 2016</u>; <u>Shabel et al., 2014</u>; <u>Wallace et al., 2017</u>). GPh-projecting dMSNs were primarily located in patches and expressed MORs (<u>Fujiyama et al., 2011</u>; <u>Stephenson-Jones et al., 2013</u>; <u>Stephenson-Jones et al., 2016</u>). This patch dMSN→GPh pathway is critical because the GPh to LHb pathway establishes the non-motor output of the basal ganglia circuitry; this contributes to the intricate evaluation of action-outcome association and action selection (<u>Fujiyama et al., 2011</u>; <u>Hong and</u> <u>Hikosaka, 2008</u>; <u>Mizumori and Baker, 2017</u>; <u>Stephenson-Jones et al., 2013</u>; <u>Stephenson-Jones et al.,</u> <u>2016</u>). These MOR-mediated striatopallidal adaptations to chronic opioid exposure and withdrawal may be essential factors driving compulsive drug-seeking and relapse.

### Chronic MOR activation by morphine elicits GABAergic striatopallidal plasticity

Chronic morphine exposure induces significant tolerance to MOR-mediated inhibition (<u>Fyfe</u> <u>et al., 2010</u>; <u>Hack et al., 2003</u>; <u>Sim-Selley et al., 2000</u>; <u>Turchan et al., 1999</u>; <u>Williams et al., 2001</u>). Tolerance is associated with a loss of functional membrane MORs due to receptor internalization (<u>Williams et al., 2013</u>). Chronic morphine exposure induces MOR redistribution from the dendritic surface to the cytoplasmic compartment in peripheral adrenergic neurons and in MSNs (<u>Drake et</u> <u>al., 2005</u>; <u>Haberstock-Debic et al., 2005</u>; <u>Haberstock-Debic et al., 2003</u>; <u>Yu et al., 2009</u>). Notably, chronic morphine treatment potentiated DAMGO-induced inhibitory currents from GABAergic interneurons onto dopaminergic neurons in the ventral tegmental area (<u>Madhavan et al., 2010</u>). We found that chronic opioid exposure induced a similar potentiation in the direct pathway. Repeated morphine exposure and withdrawal enhanced oIPSCs in GPh neurons. By strengthening GABAergic inputs to the non-motor output pathway of the basal ganglia, chronic morphine exposure could suppress GPh activity and subsequently release midbrain dopaminergic neurons from tonic GABAergic inhibition (<u>Mollick et al., 2020</u>; <u>Stephenson-Jones et al., 2016</u>). In the actor-critic model (<u>Samejima et al., 2005</u>), the patch serving as the critic will provide essential information about the state value after an action has been completed. A hyperactive patch dMSN→GPh pathway may disturb this evaluation process and assign the drug-taking behavior a greater state value; this could drive a persistent reinforcement of the behavior, leading to compulsive drug-seeking and -taking.

### Fentanyl IVSA also elicits GABAergic striatopallidal plasticity

Understanding the commonalities between natural and synthetic opioids in relation to their effects on basal ganglia circuitry would provide valuable insights into the circuit-based framework of opioid use disorder. In agreement with previous reports (<u>Bryant et al., 2021</u>; <u>Bryant et al., 2009</u>; <u>Varshneya et al., 2021</u>), we found that fentanyl induced strong hyperlocomotion and locomotor sensitization in mice, suggesting that fentanyl activates the mesolimbic dopamine pathway (<u>Di</u> <u>Chiara and North, 1992</u>). This hyperlocomotion and locomotor sensitization appear to contrast with the acute MOR-mediated suppression of striatopallidal transmission. This may be because patches account for less than 15% of the total striatal volume and do not contribute to the motor outputs (<u>Gerfen and Surmeier, 2011</u>). Fentanyl induces strong tolerance and MOR internalization (<u>Bot et al., 1998</u>; <u>Heusler et al., 2016</u>; <u>Kovoor et al., 1998</u>; <u>Terman et al., 2004</u>; <u>Virk and Williams, 2008</u>).

We found that contingent fentanyl IVSA induced similar striatopallidal adaptations as non-contingent morphine exposure. Fentanyl IVSA potentiated dMSN→GPh transmission. These data further confirmed our findings following morphine injections and suggested that both non-contingent opioid exposure and contingent opioid reinforcement potentiated dMSN outputs to the GPh.

### Fentanyl IVSA triggers GABAergic plasticity in the patch, but not matrix, compartment

Although both are within the striatum, the patch and matrix compartments have distinct neurochemical features, and local axon collaterals originating from one compartment only innervate neurons within the same compartment (<u>Banghart et al., 2015a</u>; <u>Bolam et al., 1988</u>; <u>Brimblecombe and Cragg, 2017</u>; <u>Lopez-Huerta et al., 2016</u>; <u>Prager and Plotkin, 2019</u>). Consistent with a previous finding (<u>Miura et al., 2008</u>), we showed that DAMGO suppressed IPSCs in the patch compartment, but not in the matrix compartment. We observed that fentanyl IVSA potentiated both oIPSCs and compound currents in the patches. This potentiation was accompanied by a decreased PPR and increased aIPSC frequency, indicating a presynaptic mechanism (<u>Choi and</u> <u>Lovinger, 1997</u>; <u>Dobrunz and Stevens, 1997</u>). Although most MOR expression is observed within the patch compartment, some matrix neurons are MOR^+^ (Fig. 4A) (<u>Smith et al., 2016a</u>); the physiological properties of these might resemble those of patch neurons. Accordingly, we found that only 13% of neurons could directly respond to light stimulation (< 2 ms) delivered to the matrix. Surprisingly, following chronic fentanyl exposure, these MOR^+^ matrix neurons did not show similar adaptations as patch neurons. It has been reported that both dMSNs and iMSNs in the patch send collaterals to innervate dMSNs; however, a MOR agonist exerts a greater influence on dMSN→dMSN transmission than on iMSN◊dMSN transmission (<u>Banghart et al., 2015a</u>).

Because the patch contains a significantly higher proportion of dMSNs, it is possible that most of our recorded transmission within the patch reflected dMSN→dMSN and these dMSN axons may share collaterals with dMSN→GPh projections. Both dMSN◊dMSN and dMSN◊GPh data support the view that patch dMSNs undergo profound adaptation to become hyperactive after chronic fentanyl exposure.

### Fentanyl IVSA triggers GABAergic striatonigral plasticity, a hypodopaminergic state, and negative emotional states

Numerous studies have reported that within the striatum, patch dMSNs predominantly send direct projections to SNc dopaminergic neurons (<u>Crittenden and Graybiel, 2016</u>; <u>Crittenden et</u> <u>al., 2016</u>; <u>Evans et al., 2020</u>; <u>Fujiyama et al., 2011</u>; <u>Fujiyama et al., 2015</u>; <u>McGregor et al., 2019</u>; <u>Nadel et</u> <u>al., 2021</u>; <u>Prager and Plotkin, 2019</u>). Dopamine systems play vital roles during both the initial positive reinforcement of drug-seeking and the subsequent negative reinforcement driven by hyperkatifeia during withdrawal from chronic use (<u>Koob and Volkow, 2016</u>; <u>Luscher and Malenka, 2011</u>). The emergence of hyperkatifeia after repeated opioid exposure is linked to a reduction in dopaminergic function (<u>Diana, 2011</u>; <u>Koob and Volkow, 2010</u>). Withdrawal from chronic morphine exposure decreases dopamine release in the nucleus accumbens and dorsal striatum (<u>Ashok et</u> <u>al., 2017</u>; <u>Pothos et al., 1991</u>; <u>Salinas et al., 2021</u>). We found that fentanyl IVSA reduced the spontaneous firing of SNc dopaminergic neurons. This reduction may be due to an enhancement of GABAergic signaling within the midbrain (<u>Madhavan et al., 2010</u>). We found that fentanyl IVSA potentiated dMSN→SNc GABAergic transmission, an effect that is likely to reduce the spontaneous firing of SNc dopaminergic neurons. Although high levels of dopamine D1 receptor expression have been identified in the EP and SNr/SNc (<u>Hall et al., 1994</u>), D1 receptor mRNA was only found in the striatum (<u>Beaulieu and Gainetdinov, 2011</u>; <u>Mansour et al., 1992</u>; <u>Missale et al., 1998</u>); this suggests that striatal dMSNs contain D1 receptors, while downstream neurons within the direct pathway and GABAergic interneurons in the midbrain do not express this receptor. This indicated that the ChR2-induced current mainly arose from striatal dMSNs. This result extends the above findings by demonstrating that in addition to inducing changes in dMSN projections to the EP and GPh, chronic fentanyl exposure also affected SNc dopaminergic neurons.

This hypodopaminergic state likely contributed to hyperkatifeia, which drives the negative reinforcement that propels relapse (<u>Koob, 2021</u>). Indeed, after withdrawal from repeated fentanyl injections, animals developed depression-like symptoms that were detected using multiple behavioral tasks. The emergence of depression during fentanyl withdrawal likely motivates subsequent drug use; drug seeking is reinstated after long-term withdrawal in response to contextual cues and in an effort to alleviate the depressive state (<u>Koob et al., 2020</u>). Therefore, our results indicate that a novel circuit induces hyperkatifeia following withdrawal from chronic opioid use, thus driving relapse. However, it is still intriguing to note that via the patch dMSN◊GPh◊LHb◊RMTg◊SNc pathway (<u>Stephenson-Jones et al., 2016</u>), chronic opioid-induced changes will inhibit RMTg activity; this, in turn, will activate SNc dopaminergic neurons and thus induce the opposite effect to the patch dMSN→SNc pathway.

In summary, we have discovered that chronic morphine or fentanyl exposure triggers GABAergic plasticity within striatal patches. The resultant effects on the striatopallidal and striatonigral pathways induce a hypodopaminergic state and depression-like behaviors. These results provide valuable insights into the mechanisms underlying the opioid-induced negative emotional states that drive opioid relapse.

## STAR METHODS

### Reagents

TTX, CTAP, and DAMGO were obtained from Tocris. RetroBeads were obtained from LumaFluor. Morphine sulfate pentahydrate was a gift from the National Institute on Drug Abuse Drug Supply Program. Fentanyl citrate (Cat. #1270005), 4-aminopyridine (4-AP), picrotoxin, and all other chemicals were purchased from Sigma-Aldrich.

### Key resources table

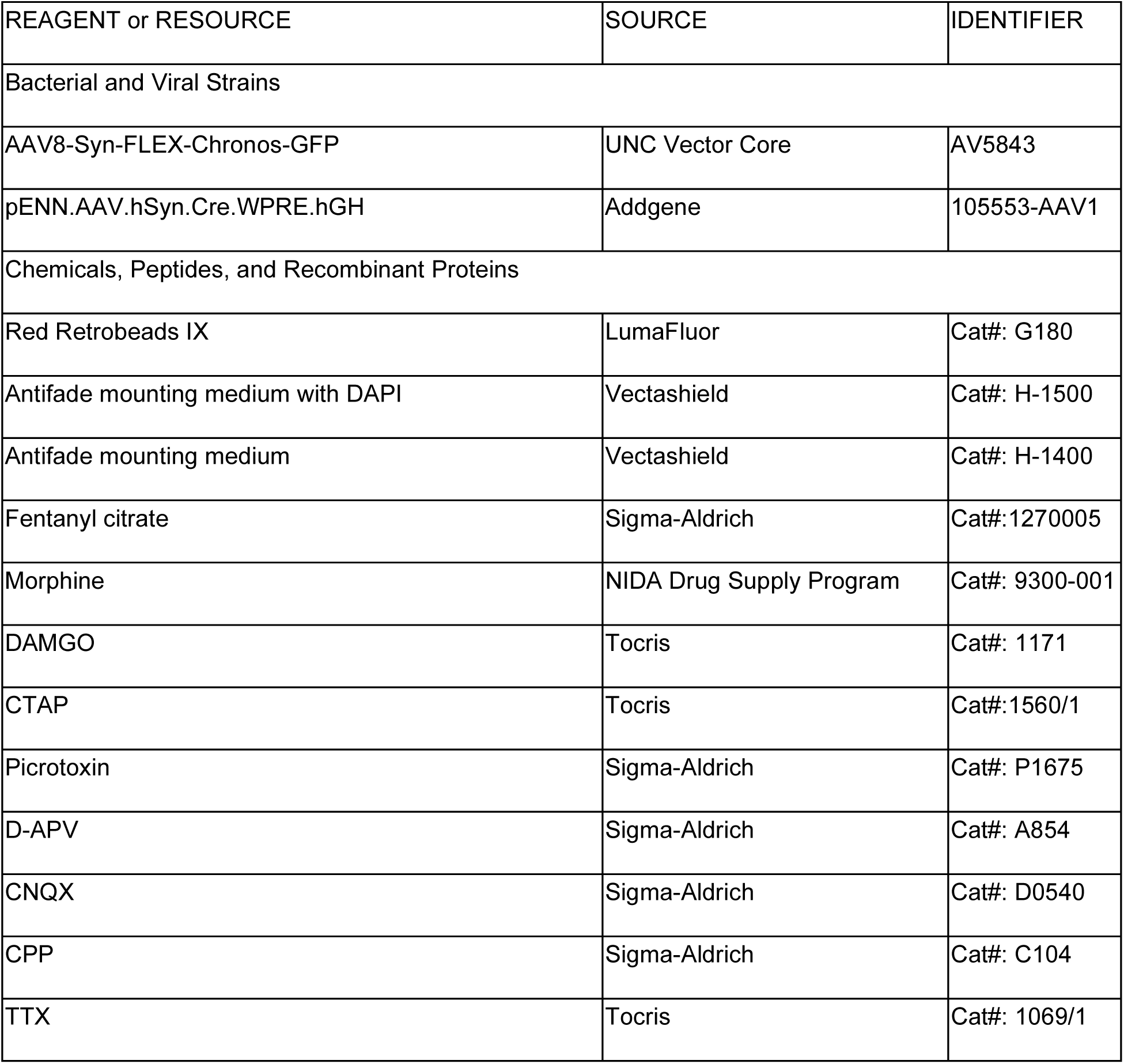
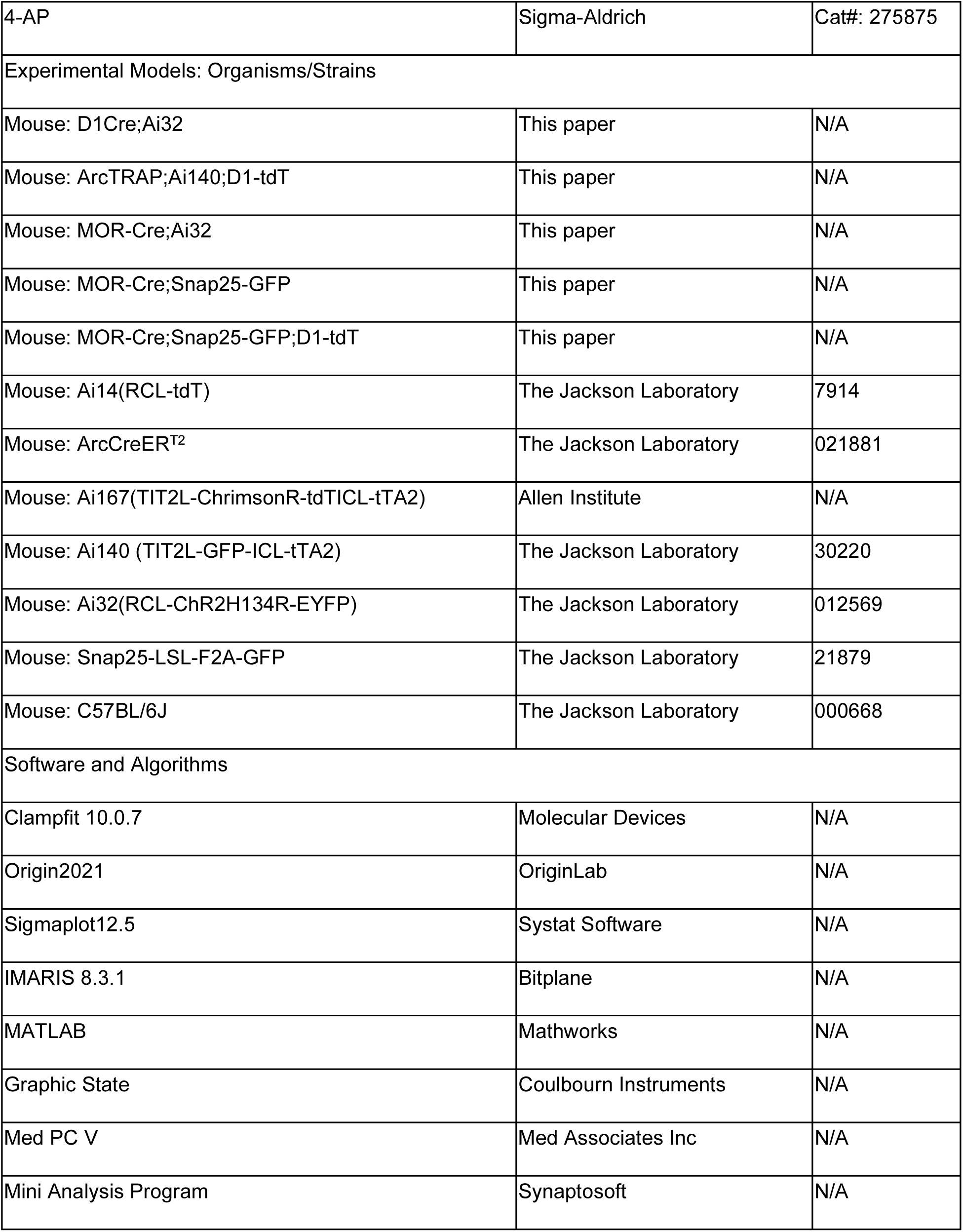

### Animals

The *Drd1a*-Cre (D1-Cre) mice, MOR-Cre mice, and Long Evans rats were obtained from Mutant Mouse Regional Resource Centers. D1-tdTomato, ArcCreER^T2^ (ArcTRAP), Ai167(TIT2L- ChrimsonR-tdTICL-tTA2), Ai140 (TIT2L-GFP-ICL-tTA2), Ai32(RCL-ChR2H134R-EYFP), Ai14(RCL-tdT), Snap25-LSL-F2A-GFP, wild-type (WT) C57BL/6J mice were purchased from the Jackson Laboratory. D1-Cre or MOR-Cre mice were crossed with Ai32 to generate D1-Cre;Ai32 or MOR-Cre;Ai32 mice, respectively. D1-Cre mice (<u>Bailly et al., 2020</u>) were crossed with Ai167 mice to generate D1-Cre;Ai167 mice, and ArcTRAP mice were crossed with Ai140 and D1-tdT mice to generate the triple transgenic ArcTRAP;Ai140;D1-tdT mice. Genotyping was performed using PCR of tail DNA. Both male and female transgenic littermates (3–4 months old) were used in this study. Animals were housed with 2–4 gender-matched conspecifics in a 12-h light-dark cycle (light from 7 a.m. to 7 p.m.) with *ad libitum* food and water. The room temperature for all animal housing and experiments was 23°C. Morphine and fentanyl were diluted with 0.9% saline before use. Mice were treated with i.p. morphine injections at escalating doses of 20, 40, 60, 80, and 100 mg/kg once daily for 5 consecutive days, or with the same volume of 0.9% saline. Mice received 7 daily injections of saline or fentanyl (0.3 mg/kg, i.p). Mice with different genotypes were randomly assigned to morphine-, fentanyl- or saline-treated groups. Investigators were blind to all experimental groups. All animal care and experimental procedures were approved by the Texas A&M University Institutional Animal Care and Use Committee and were conducted in agreement with the National Research Council Guide for the Care and Use of Laboratory Animals.

### Electrophysiological recordings of brain slices

#### Brain slice preparation

Slices were prepared as previously described (<u>Lu et al., 2019</u>; <u>Ma</u> <u>et al., 2018</u>; <u>Ma et al., 2021</u>). Mice were anesthetized with isoflurane and subsequently decapitated. The brains were removed rapidly and sagittal slices of the striatum with the EP (230 µm in thickness) or coronal slices (250 µm in thickness) of the striatum or SNc were cut in an ice-cold cutting solution containing the following (in mM): 40 NaCl, 143.5 sucrose, 4 KCl, 1.25 NaH_2_PO_4_, 26 NaHCO_3_, 0.5 CaCl_2_, 7 MgCl_2_, 10 glucose, 1 sodium ascorbate and 3 sodium pyruvate (pH 7.35, 305–310 mOsm), saturated with 95% O_2_ and 5% CO_2_. Slices were then incubated in a 1:1 mixture of cutting solution and external solution at 32°C for 45 min. The external solution was composed of the following (in mM): 125 NaCl, 4.5 KCl, 2 CaCl_2_, 1 MgCl_2_, 1.25 NaH_2_PO_4_, 25 NaHCO_3_, 15 sucrose and 15 glucose, pH 7.35, 305–310 mOsm), saturated with 95% O_2_ and 5% CO_2_. Slices were maintained in external solution at room temperature until use.

#### Electrophysiological recordings

Recordings were performed as previously described (<u>Cheng et al., 2018</u>; <u>Huang et al., 2017</u>; <u>Lu et al., 2019</u>; <u>Ma et al., 2018</u>; <u>Ma et al., 2021</u>). The brain slice was positioned in a recording chamber attached to the fixed stage of an upright microscope (Olympus) and perfused with oxygenated external solution at 32°C, with a flow rate of 2 mL/min. Neurons were visualized using a 40× water-immersion lens and an infrared-sensitive CCD camera. A Multiclamp 700B amplifier with Clampex 10.6 software and Digidata1550A data acquisition system (Molecular Devices, Sunnyvale, CA) were used for recording. A recording pipette with a resistance of 3–6 MΩ was pulled from a borosilicate glass capillary (World Precision Instruments, Sarasota, FL) using a micropipette puller (Model P-97, Sutter Instrument Co.).

Whole-cell voltage-clamp recordings were used to measure GABAergic synaptic currents in striatal MSNs, GPh, and SNc dopaminergic neurons. Recording electrodes contained internal solution (in mM: 125 CsMeSO4, 6 NaCl, 10 HEPES, 1 EGTA, 10 QX-314.Cl, 2 MgATP, 6 Na_3_GTP, 2 Na_2_CrPO_4_; pH 7.25, 280 mOsm). DNQX (20 µM) and D-AP5 (50 µM) were included in the external solution. Neurons were clamped at -70 mV (MSNs or GPh neurons) or -60 mV (SNc dopaminergic neurons). To measure electrically evoked IPSCs in patch and matrix neurons, a bipolar glass stimulating electrode was placed in the same compartment, around 100 µm away from the neuron that was being recorded; a brief pulse (40 µA) was delivered at 0.05 Hz. To measure the input-output responses of optogenetically-induced GABAR-IPSCs in MSNs, GPh neurons, and SNc dopaminergic neurons, stimulations were conducted using a series of light intensities delivered at 470 nm (Chronos or ChR2) or 590 nm (Chrimson) for 2 ms. For some spatially specific experiments, picrotoxin (100 µM), TTX (1 µM), and/or 4-AP (500 µM), were added to block GABA_A_R-mediated currents and voltage-gated sodium and potassium channels, respectively. DAMGO (0.3 or 1 µM) or CTAP (1 µM) were perfused for 5–10 min to activate or inhibit MORs, respectively. To measure the probability of transmitter release, we delivered a pair of light stimulations at an interval of 100 ms. The PPR was calculated by dividing the amplitude of the second IPSC by the amplitude of the first IPSC. To measure aIPSCs, the Ca^2+^ in the external solution was replaced by Sr^2+^ (0.5 mM) (<u>Cheng et al.,</u> <u>2021</u>), and the aIPSC frequency and amplitudes were collected from 100–600 ms after each stimulus. The frequency and amplitudes of sIPSCs were recorded under gap-free conditions for 3 min. *I*_h_ currents in SNc dopaminergic neurons were detected under whole-cell voltage-clamp mode (hold potential: -60 mV) using a series of voltage steps from -150 mV to 0 mV (1 sec, 30- mV increments) using the same internal and external solutions that were used for action potential recording (<u>Tan et al., 2021</u>).

Cell-attached voltage-clamp or whole-cell current-clamp techniques were performed to record spontaneous action potentials of SNc dopaminergic neurons, without antagonists in the external solution. The pipette was filled with a K^+^-based intracellular solution (in mM: 123 potassium gluconate, 10 HEPES, 0.2 EGTA, 8 NaCl, 2 MgATP, and 0.3 NaGTP; pH 7.25, 280 mOsm). Unlike whole-cell patch-clamp recording, which clamped neurons close to -60mV, cell-attached patch-clamping recorded the firing of neurons without a holding potential, when the resistance exceeded 1 GΩ. Light activation was achieved using a blue (470-nm) LED (DC4104, Thorlabs) delivered at 1–20 Hz with a 2-ms duration (10 pulses). Evoked action potentials or *I* ^-^ mediated sag were measured using whole-cell current-clamping with increasing current injections (from 0–120 pA or from -120–0 pA, in 1-sec incremental step). The sag amplitudes were calculated by subtracting the steady-state voltage at the end of each current step from the trough potential at the beginning of each current step.

### Surgery

#### Stereotaxic virus infusion

Virus infusion was conducted as described previously (<u>Cheng et</u> <u>al., 2021</u>; <u>Lu et al., 2021b</u>; <u>Ma et al., 2021</u>). Animals were anesthetized with 3–4% isoflurane at 1.0 L/min and immobilized in a stereotaxic surgery frame with ear bars (David Kopf Instruments); lubricant ophthalmic ointment was applied to prevent eye drying. After a midline scalp incision, small bilateral craniotomies were made using a microdrill (0.5 mm burr), based on the mouse brain atlas. AAV-Flex-Chronos-GFP and AAV1-Cre were infused into the DMS (AP1: +0.38 mm, ML1: ±1.55 mm, DV1: -2.9 mm; AP2: +0.02 mm, ML2: ±1.8 mm, DV2: -2.7 mm from the Bregma) and red beads were infused into the LHb (AP: -1.7 mm, ML: 0.53 mm, DV: 2.8 mm from the Bregma) using glass capillaries (GASTIGHT) at a rate of 0.12 µL/min (0.3–0.5 µL/site, bilaterally). The injectors were maintained at the injection sites for 10 min after the end of the infusion to allow virus diffusion. After suturing the scalp incision, the animals were returned to their home cages for at least 6 weeks for recovery and virus expression.

#### Jugular vein catheter implantation

The jugular vein catheter implantation was performed as described previously (<u>Thomsen et al., 2005</u>). Briefly, each mouse was anesthetized with an i.p. injection of ketamine (120 mg/kg) and xylazine (6 mg/kg), followed by subcutaneous administration of ketoprofen (0.05 mL of a 1-mg/mL solution). Two incision sites were made at the dorsal mid-scapular region and the ventral upper-right torso following disinfection by scrubbing with a 70% alcohol pad. A vascular access button (Instech, VABM1BSM/25) connected to a jugular vein catheter (Instech, C20PU-MJV1934) was then placed subcutaneously over the right shoulder, and the catheter was inserted 1.2-cm into the right jugular vein under sterile conditions. Two suture threads were tied above and below the anchoring beads around the vein to secure the intracardial tubing. After surgery, mice were allowed a 7-day recovery period, during which 0.02 mL saline containing heparin (30 USP units/mL) and cefazolin (67 mg/mL) was administered daily via the catheter to prevent clotting and infection. The free end of the cannula guide was kept closed before and after experiments. Catheter patency was confirmed by a loss of righting reflex within 5 sec of infusing 0.03 mL of ketamine (15 mg/mL) plus midazolam (0.75 mg/mL) in saline through the catheter. Patency was checked weekly, and animals with clogged catheters were removed from the study.

### Behavioral testing

***Open field test*** was conducted as we previously described (<u>Cheng et al., 2018</u>; <u>Hellard et al.,</u> <u>2019</u>; <u>Huang et al., 2017</u>). A transparent open field activity chamber (Med Associates, 43 cm × 43 cm × 21 cm height) was equipped with an infrared beam detector that was connected to a computer. Animals were placed into the chamber 15 min after i.p. injections of saline, morphine (20, 40, 60, 80, 100 mg/kg) (<u>Varshneya et al., 2021</u>) or fentanyl (0.3 mg/kg) (<u>Bryant et al., 2021</u>; <u>Bryant</u> <u>et al., 2009</u>), and activity was monitored for 30 min. The distance traveled, velocity, and time spent on the periphery were analyzed by Activity Monitor software (MED Associates, St Albans, VT). Heatmaps were generated using scripts in MATLAB (MathWorks). For analysis of fentanyl- withdrawal induced negative emotional states, animals were habituated in the chamber for 10 min one week before the open field test, and locomotor activity was measured for 10 min on day 10 of fentanyl withdrawal. The immobility time and distance traveled were analyzed by Activity Monitor software.

#### IVSA of fentanyl

After the 7-day recovery from implantation surgery, mice were trained in operant chambers (Med Associates, St Albans, VT) housed in a light- and sound-attenuating chamber. The operant chambers contained two nose-poke holes, with cue lights inside and above. To deliver fentanyl or saline, a cannula running through the ceiling of the chamber could be attached to the jugular vein catheter. The cue light above the active nose-poke hole was illuminated at the beginning of each session, indicating fentanyl availability, and extinguished at the end of the session. Nose pokes in the active hole turned off this cue light and triggered a 3.2- sec fentanyl (3 µg/kg) infusion (<u>Criado and Gomez e Segura, 2003</u>; <u>Muelbl et al., 2016</u>), which coincided with a 3.2-sec audio cue and a cue light in the active hole (<u>Van den Oever et al., 2008</u>).

After fentanyl delivery, the light above the active hole remained off for 20 sec (time-out period). During the time-out period, responses in the nose-poke ports were recorded but did not result in fentanyl delivery. Mice were trained to self-administer fentanyl using daily 3-h sessions (5 d/week) under a fixed ratio 1 (FR1) schedule. Once responses had stabilized under the FR1 schedule and met defined criteria (> 10 infusions with > 75% responses at the active port; < 20% variation in 3 consecutive sessions) (<u>Ezeomah et al., 2020</u>; <u>Muelbl et al., 2016</u>; <u>Stevenson et al., 2020</u>; <u>Sustkova-Fiserova et al., 2020</u>), animals were moved to FR2 and then FR3 schedules.

#### Social interaction test

Social interactions were investigated in a white acrylic chamber (45 cm wide × 45 cm long × 45 cm high). A stranger mouse was placed in a metal cage (8 cm in diameter × 11 cm high) within a 16-cm diameter “interaction” zone along one wall of the chamber. “Avoidance” zones (10 cm × 10 cm) were designated in both opposite corners of the open field. The test mouse was allowed to explore the chamber freely and its behavior was tracked using an overhead camera and video-tracking software (Ethovision XT, Noldus). The time spent in the “interaction” and “avoidance” zones and the number of entries into each were analyzed.

#### Sucrose splash test

Immediately after social interaction testing, sucrose splash testing was conducted over a 5-min trial in the white acrylic chamber (45 cm wide × 45 cm long × 45 cm high). The test mouse was placed in the chamber and a 10% sucrose solution was pipetted onto its dorsal coat. Behavior was recorded using an overhead camera and scored by a treatment-blinded experimenter. The percentage of total trial time spent grooming was determined, and the mouse was then returned to its home cage.

#### Conditioned place preference (CPP)

CPP tests were conducted as we prevoulsy described (<u>Cheng et al., 2017a</u>; <u>Cheng et al., 2018</u>). A two-compartment CPP apparatus (Med Associates) was used, illuminated by 45 lux. The two compartments had different floor textures (bar or grid), wall colors (black or white), and bedding textures (strips or pellets). They were separated by a manual guillotine door and were equipped with infrared photobeams connected to a computer; these recorded the animals’ locations. The procedure consisted of pre-conditioning (day 0), conditioning (days 1–7), and post-conditioning (days 8 and 36) tests. On the pre- conditioning day, all the rats were i.p. injected with saline and allowed to freely explore both compartments for 15 min, with the middle door open. During the 30-min conditioning sessions, the manual guillotine door was closed to restrict animals to one compartment. In the fentanyl group, half of the rats received fentanyl (0.03 mg/kg, i.p.) (<u>Gaulden et al., 2021</u>) in the morning in one compartment and saline (i.p.) in the afternoon in the other compartment, while the other half received saline in the morning and fentanyl in the afternoon. In the saline group, all rats received saline during both morning and afternoon sessions. We repeated this process of contextual pairing of fentanyl and saline administration for a total of 7 conditioning days. On the post- conditioning days, rats were returned to the compartments for 15 min with the middle gate open; this allowed us to measure CPP after drug withdrawal for 1 day and 28 days. The CPP score was calculated by subtracting the time spent in the non-paired compartment from the time spent in the fentanyl-paired compartment.

### Histology and cell counting

Histology studies were conducted as we previously described (<u>Lu et al., 2021a</u>; <u>Wei et al., 2018</u>). Mice were perfused with PBS and fixed with 4% paraformaldehyde. Brains were removed and post-fixed in 4% paraformaldehyde for 24 h, dehydrated with 30% sucrose for 3 d at 4°C, and coronally cut into 50-μm slices at -22°C. After washing in PBS, slices were mounted on slides with VECTASHIELD mounting media (Vector Laboratories). Images were acquired by confocal microscopy (Fluoroview-1200, Olympus) and analyzed by Imaris 8.3.1 (Bitplace, Zurich, Switzerland). All specimens were imaged and analyzed using identical parameters.

### Cluster analysis based on t-distributed stochastic neighbor embedding (tSNE)

tSNE cluster analysis was performed using MATLAB as described previously (<u>Li et al., 2021b</u>). The variables included were derived from the behavioral tests conducted 10 d after the last fentanyl injection; these included locomotor activity testing (immobility time, total distance traveled, distance traveled over time), social interaction testing (velocity, distance moved, avoidance zone visits, proximity zone visits, time spent in avoidance/proximity zone), and sucrose splash testing (latency to first grooming, grooming frequency, time spent grooming).

### Statistical analysis

Electrophysiological data analyses were performed using the Clampfit software (Molecular Devices, Sunnyvale, CA) and Mini Analysis Program (Synaptosoft, Decatur, GA). All statistical analyses were performed by SigmaPlot 12.5 (Systat Software Inc.). Experiments with more than two groups were subjected to one-way ANOVA, two-way ANOVA, one-way repeated measures ANOVA (RM ANOVA), or two-way RM ANOVA with Turkey post-hoc tests for multiple comparisons. Experiments with two groups were analyzed using two-tailed paired or unpaired Student’s *t* tests. All data are presented as the mean ± s.e.m.

## ACKNOWLEDGEMENTS

We thank Dr. Brigitte L. Kieffer for providing us with the MOR-Cre mouse line that was utilized to generate the MOR-Cre;Ai32, MOR-Cre;Snap25-GFP, and MOR-Cre;Snap25-GFP;D1-tdT mice, which we used to study on neurons expressing the mu opioid receptor. We thank Sebastian Melo and Jared Jarger for technical assistance, and John Chainaranont, Joanna Mendoza, and Matthew J. Childs for their help in data analysis. This work was supported by NIH grants: U01AA025932 (to JW), R01AA021505 (to JW), R01AA027768 (to JW); and by an X-Grant from the Presidential Excellence Fund at Texas A&M University (to JW).

## AUTHOR CONTRIBUTIONS

JW conceived, designed, and supervised all the experiments in the study. WW, XX, and JW wrote the first draft of the manuscript. WW, XZ, HG, and ZH designed and performed electrophysiological experiments and analyzed the data. XX, XZ, YFH, TT, AS, MH, and ED designed behavioral experiments. XX, XZ, YFH, TT, AS, and RC performed the behavioral experiments and analyzed the data. WP, TT, and EY conducted histology experiments and analyzed the data. DW bred transgenic animals and conducted genotyping. JW, MH, YH, and ED discussed and revised the manuscript. WW and XX contributed equally to this research as co-first authors. The order of co-first authors was determined by the temporal order of research contribution and was agreed upon by the co-first authors. Co-first authors have the right to list themselves first for purposes of their curriculum vitae and biosketch.

## DECLARATION OF INTERESTS

The authors declare no competing interests.

## SUPPLEMENTARY FIGURE LEGENDS

**Supplementary Figure 1.**
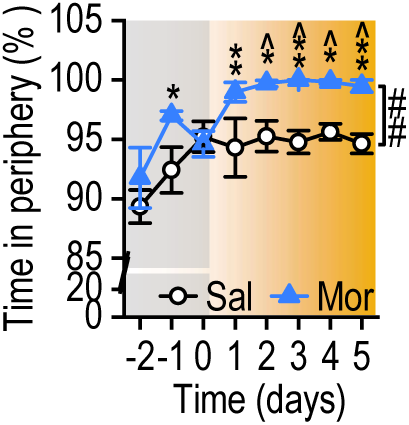
Repeated morphine administration increases peripheral locomotion. Time-course of time spent in the periphery for saline- and morphine-treated mice. ^##^*p* < 0.01 by two-way RM ANOVA; \**p* < 0.05, \*\**p* < 0.01 versus the same day in the saline group; *^p* < 0.05 versus day 0 within the morphine group by Tukey *post hoc* test; n = 4 (Sal) and 6 (Mor) mice.

**Supplementary Figure 2.**
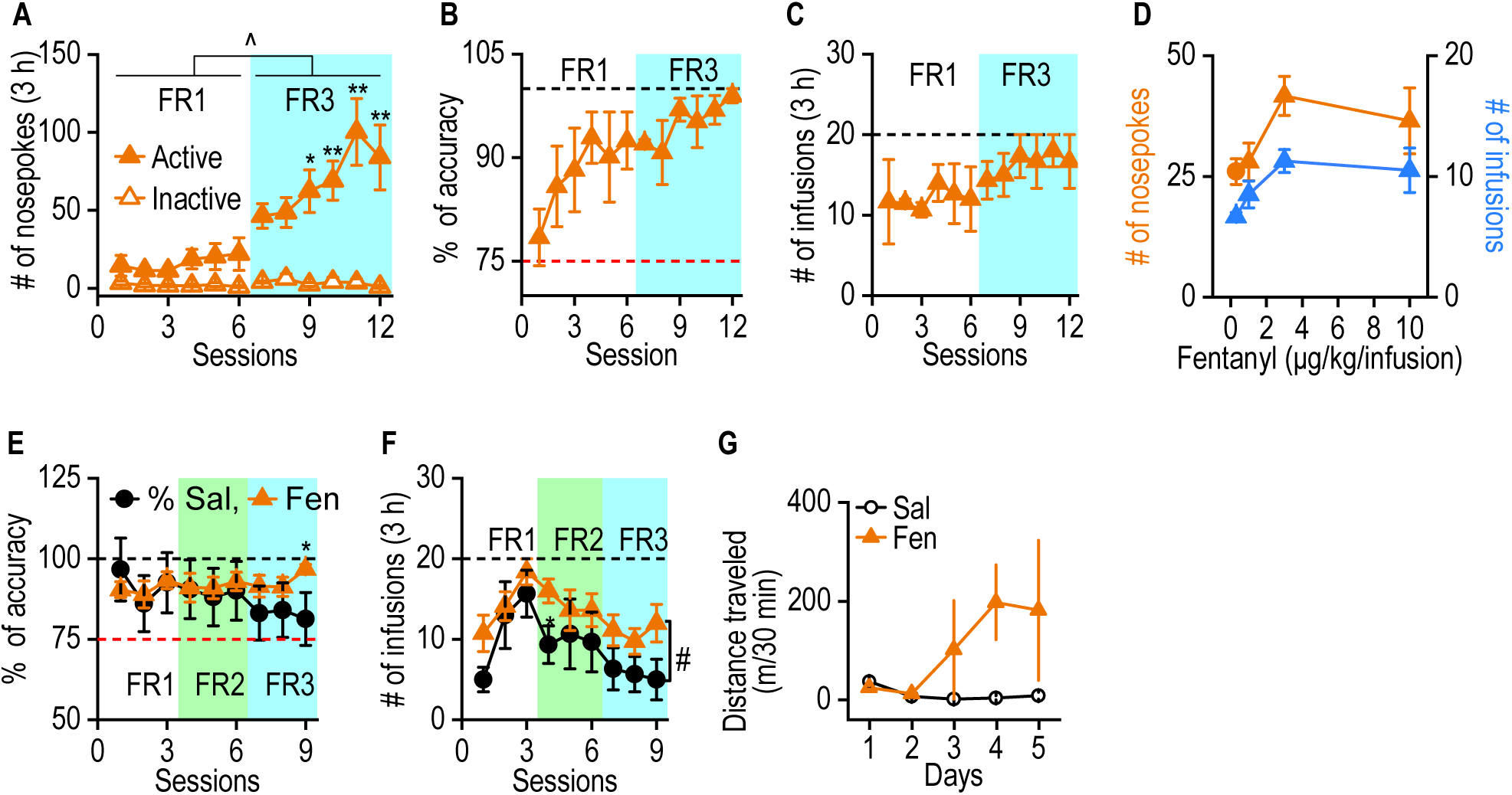
Intravenous self-administration of fentanyl. C57BL/6 (*A–D*) or MOR-Cre (D1-Cre);Ai32 (*E*, *F*) mice were trained to self-administer fentanyl (3 µg/kg/infusion) or saline (0.9%) using FR1–3 schedules. ***A***, Fentanyl-induced effects on nosepokes from FR1 to FR3; ^^^*p* < 0.05 for FR1 versus FR3, for both active and inactive nosepokes; \**p* < 0.05, \*\**p* < 0.01 for the comparison of active and inactive nosepokes during the same session, Tukey *post hoc* test. ***B***, Accuracy (active nosepokes expressed as a percentage of total nosepokes) for fentanyl was persistently high and was slightly increased after training; *p* > 0.05. An accuracy of 75% was achieved before proceeding to the next FR schedule. ***C***, Animals achieved a consistent number of infusions. The maximum number of infusions was 20; *p* > 0.05 for FR1 versus FR3. ***D***, Inverted U-shaped dose-response curves were observed for active nosepokes and fentanyl infusions. ***E***, Accuracy of active nosepokes for mice receiving fentanyl or saline IVSA; \**p* < 0.05, Tukey *post hoc* test. ***F***, Number of infusions earned by fentanyl and saline IVSA mice. A maximum of 20 infusions were delivered during each session. ^#^*p* < 0.05 for saline versus fentanyl active presses; \**p* < 0.05 versus the saline group during the same session, Tukey *post hoc* test. ***G***, Time-course of i.p. fentanyl (Fen, 0.3 mg/kg)-induced hyperactivity in ArcTRAP;Ai140;D1-tdT mice. Two-way RM ANOVA for *A* and *E*–*G*, and one-way RM ANOVA for *B*–*D*; n = 3 (*A*–*D*), 8 (*E*, *F*; fentanyl), 3 (*E*, *F*; saline), and 2 per group (*G*).

**Supplementary Figure 3.**
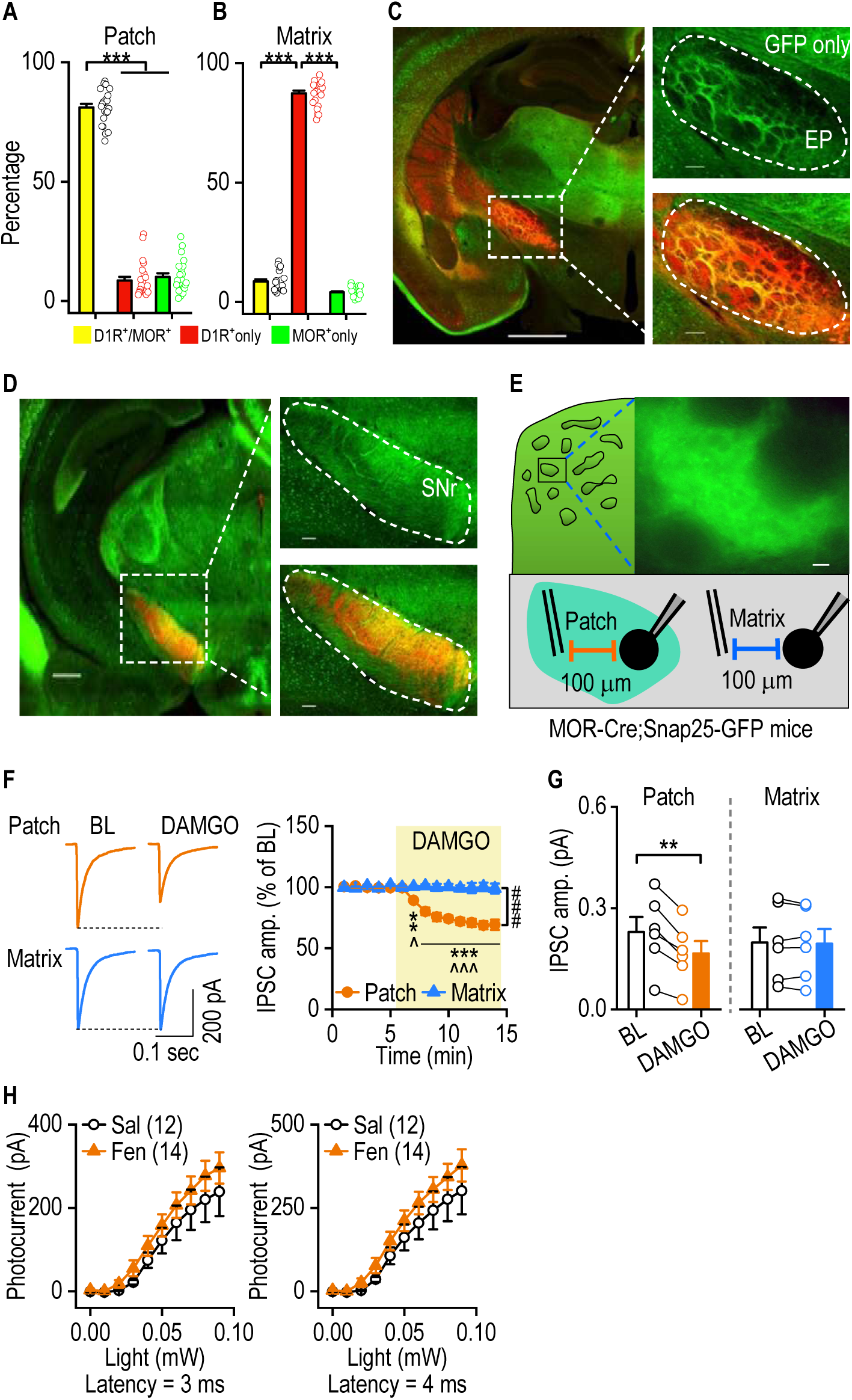
Acute and chronic MOR activation exert opposite effects on GABAergic transmission in DMS patch neurons. ***A, B***, The percentage of MOR-expressing dMSNs (GFP^+^ and tdT^+^) was much higher in the patch compartment (*A*) than in the matrix compartment (*B*); ***p < 0.001. ***C, D***, Representative images of sagittal sections from a MOR-Cre;Snap25-GFP;D1-tdT mouse showing MOR- and GFP- expressing fibers in the EP (*C*) and SNr (*D*) areas. Scale bar, 1 mm (*C*, left), 0.1 mm (*C*, right; *D*, right), and 0.5 mm (*D*, left). ***E***, Schematic showing the stimulating and recording electrodes (placed 100 μm apart) within the patch and matrix compartments of MOR-Cre;Snap25-GFP mice. ***F, G***, Bath application of DAMGO inhibited electrically evoked IPSCs in patch, but not matrix, neurons. Time-course (*F*): ^###^*p* < 0.001; \*\**p* < 0.01, \*\*\**p* < 0.001 versus the matrix group at the same timepoint; ^*p* < 0.05, ^^^*p* < 0.001 versus the baseline in the patch group. Bar graph (*G*) for baseline (BL; 5-min average) versus DAMGO (14th min) in patch (left) and matrix (right) compartments; \*\**p* < 0.01. ***H***, In MOR^+^ patch neurons, fentanyl IVSA did not alter I_photo_ measured at 3 ms (left) or 4 ms (right) from the start of laser stimulation. One-way ANOVA (*A, B*), two-way RM ANOVA (*F, H*), and paired *t* test (*G*); n = 6/4 (*F, G*; patch), 6/3 (*F, G*; matrix), 12/3 (*H*; Sal), 14/3 (*H*; Fen).

**Supplementary Figure 4.**
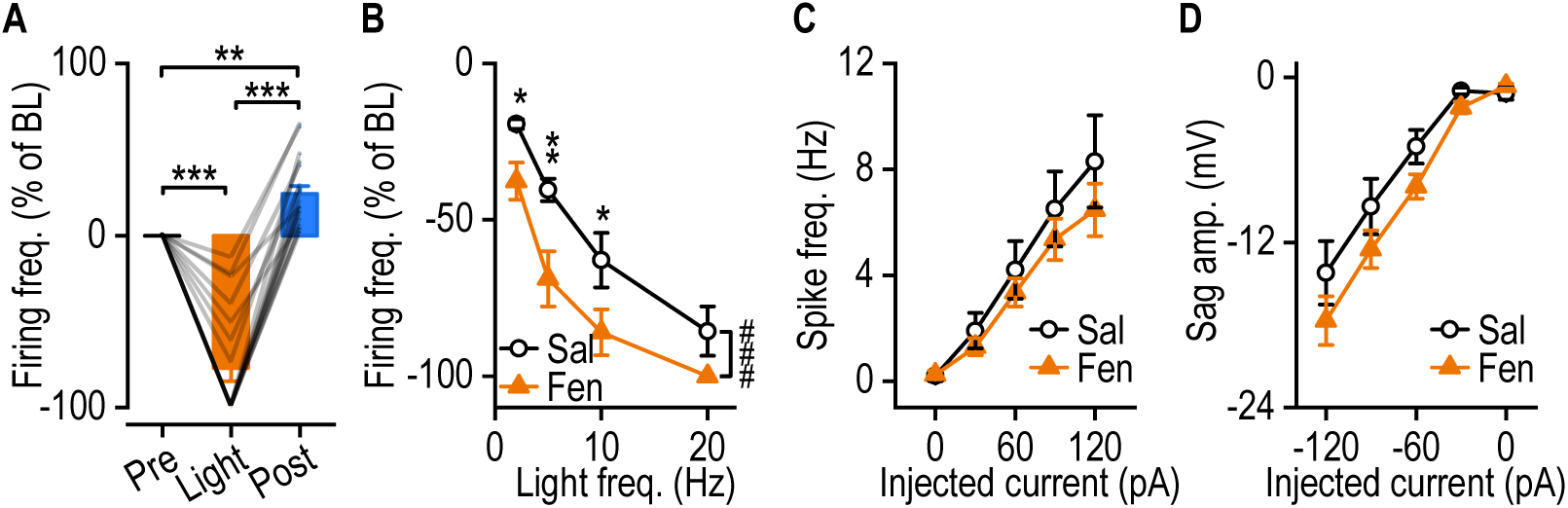
Fentanyl IVSA suppresses spontaneous firing without altering the intrinsic excitability of SNc dopaminergic neurons. ***A***, Optogenetic excitation of D1R^+^ inputs inhibited spontaneous firing and caused a rebound potentiation of firing in SNc dopaminergic neurons from D1-Cre;Ai32 mice; \*\**p* < 0.01, \*\*\**p* < 0.001. Pre, Light, and Post: before, during, and after a burst light stimulation (10 pulses, 20 Hz). ***B***, Fentanyl IVSA increased the D1R-derived suppression of spontaneous firing in SNc dopaminergic neurons; ^###^*p* < 0.001; \**p* < 0.05 and \*\**p* < 0.01, Tukey *post hoc* test. ***C, D***, Fentanyl IVSA did not alter evoked firing frequencies (*C*) or sag amplitudes (*D*); *p* > 0.05. One-way RM ANOVA (*A*), two- way RM ANOVA (*B*–*D*). n = 18/3 (*A*), 14/2 (*B*; Sal), 11/3 (*B*; Fen), 10/3 (*C*; Sal), 15/5 (*C*; Fen), 10/3 (*D*; Sal), 16/5 (*D*; Fen).

**Supplementary Figure 5.**
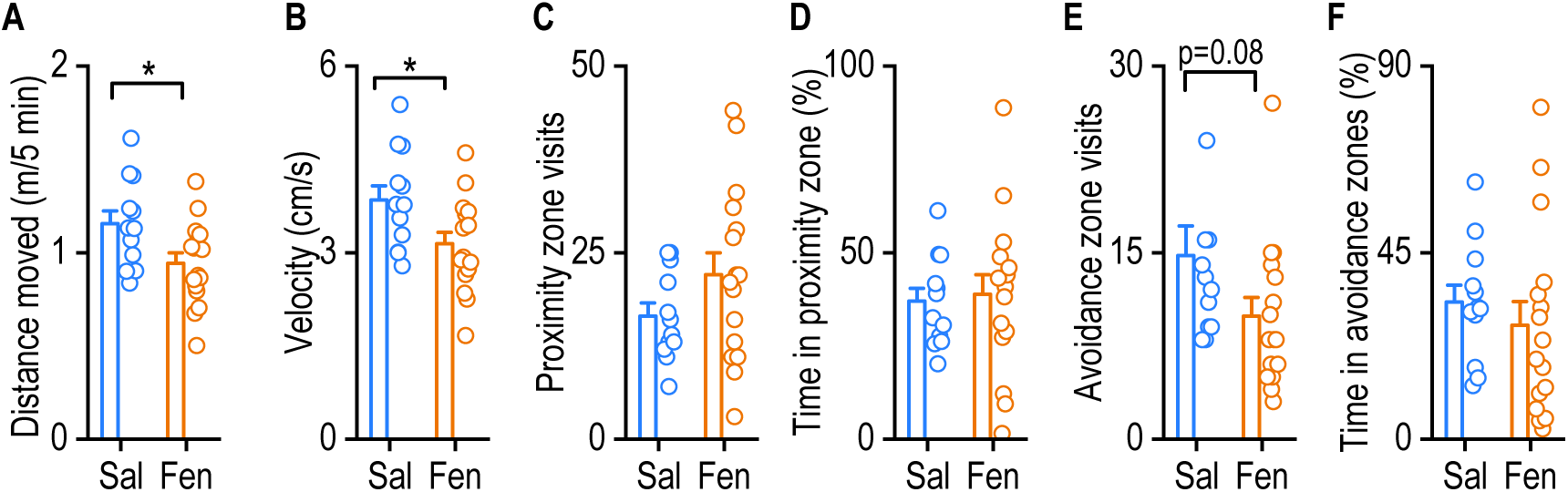
Withdrawal from repeated fentanyl injections reduces locomotor activity during social interaction testing. C57BL/6 mice received daily fentanyl injections for 14 d, and a social interaction test was conducted 10 d after the last injection. ***A***, ***B***, Fentanyl withdrawal reduced the distance traveled (*A*) and velocity (*B*) during the social interaction test; \**p* < 0.05. ***C–F***, Fentanyl withdrawal did not cause any detectable changes in the proximity zone visits (*C*), time in proximity zones (*D*), avoidance zone visits (*E*), or time in avoidance zones (*F*). Unpaired *t* test (*A*–*F*); n = 12 (Sal) and 16 (Fen).

## Notes

### Competing Interest Statement

The authors have declared no competing interest.

## REFERENCES

Al-Hasani, R., and Bruchas, M.R. (2011). Molecular mechanisms of opioid receptor-dependent signaling and behavior. Anesthesiology 115, 1363–1381.

Ashok, A.H., Mizuno, Y., Volkow, N.D., and Howes, O.D. (2017). Association of Stimulant Use With Dopaminergic Alterations in Users of Cocaine, Amphetamine, or Methamphetamine A Systematic Review and Meta-analysis. Jama Psychiat 74, 511–519.

Bailly, J., Del Rossi, N., Runtz, L., Li, J.J., Park, D., Scherrer, G., Tanti, A., Birling, M.C., Darcq, E., and Kieffer, B.L. (2020). Targeting morphine-responsive neurons: generation of a knock-in mouse line expressing Cre recombinase from the mu opioid receptor gene locus. eNeuro 7, 0433-0419.2020.

Baker, P.M., Jhou, T., Li, B., Matsumoto, M., Mizumori, S.J., Stephenson-Jones, M., and Vicentic, A. (2016). The Lateral Habenula Circuitry: Reward Processing and Cognitive Control. J Neurosci 36, 11482–11488.

Bamford, N.S., Wightman, R.M., and Sulzer, D. (2018). Dopamine’s effects on corticostriatal synapses during reward-based behaviors. Neuron 97, 494–510.

Banghart, M.R., Neufeld, S.Q., Wong, N.C., and Sabatini, B.L. (2015a). Enkephalin Disinhibits Mu Opioid Receptor-Rich Striatal Patches via Delta Opioid Receptors. Neuron 88, 1227–1239.

Banghart, M.R., Neufeld, S.Q., Wong, N.C., and Sabatini, B.L. (2015b). Enkephalin Disinhibits Mu Opioid Receptor-Rich Striatal Patches via Delta Opioid Receptors. Neuron 88, 1227–1239.

Beaulieu, J.M., and Gainetdinov, R.R. (2011). The physiology, signaling, and pharmacology of dopamine receptors. Pharmacol Rev 63, 182–217.

Bolam, J.P., Izzo, P.N., and Graybiel, A.M. (1988). Cellular substrate of the histochemically defined striosome/matrix system of the caudate nucleus: a combined Golgi and immunocytochemical study in cat and ferret. Neuroscience 24, 853–875.

Bolam, J.P., and Smith, Y. (1992). The striatum and the globus pallidus send convergent synaptic inputs onto single cells in the entopeduncular nucleus of the rat: a double anterograde labelling study combined with postembedding immunocytochemistry for GABA. J Comp Neurol 321, 456–476.

Bot, G., Blake, A.D., Li, S., and Reisine, T. (1998). Fentanyl and its analogs desensitize the cloned mu opioid receptor. J Pharmacol Exp Ther 285, 1207–1218.

Brimblecombe, K.R., and Cragg, S.J. (2017). The Striosome and Matrix Compartments of the Striatum: A Path through the Labyrinth from Neurochemistry toward Function. ACS Chem Neurosci 8, 235–242.

Bryant, C.D., Healy, A.F., Ruan, Q.T., Coehlo, M.A., Lustig, E., Yazdani, N., Luttik, K.P., Tran, T., Swancy, I., Brewin, L.W., et al. (2021). Sex-dependent effects of an Hnrnph1 mutation on fentanyl addiction-relevant behaviors but not antinociception in mice. Genes Brain Behav 20, e12711.

Bryant, C.D., Roberts, K.W., Culbertson, C.S., Le, A., Evans, C.J., and Fanselow, M.S. (2009). Pavlovian conditioning of multiple opioid-like responses in mice. Drug and alcohol dependence 103, 74–83.

Chan, K.Y., Jang, M.J., Yoo, B.B., Greenbaum, A., Ravi, N., Wu, W.L., Sanchez- Guardado, L., Lois, C., Mazmanian, S.K., Deverman, B.E., and Gradinaru, V. (2017). Engineered AAVs for efficient noninvasive gene delivery to the central and peripheral nervous systems. Nat Neurosci 20, 1172–1179.

Cheng, Y., Huang, C.C.Y., Ma, T., Wei, X., Wang, X., Lu, J., and Wang, J. (2017a). Distinct synaptic strengthening of the striatal direct and indirect pathways drives alcohol consumption. Biol Psychiatry 81, 918–929.

Cheng, Y., Huang, C.C.Y., Ma, T., Wei, X., Wang, X., Lu, J., and Wang, J. (2017b). Distinct Synaptic Strengthening of the Striatal Direct and Indirect Pathways Drives Alcohol Consumption. Biological psychiatry 81, 918–929.

Cheng, Y., Wang, X., Wei, X., Xie, X., Melo, S., Miranda, R.C., and Wang, J. (2018). Prenatal exposure to alcohol induces functional and structural plasticity in dopamine D1 receptor-expressing neurons of the dorsomedial striatum. Alcohol Clin Exp Res 42, 1493–1502.

Cheng, Y., Xie, X., Lu, J., Gangal, H., Wang, W., Melo, S., Wang, X., Jerger, J., Woodson, K., Garr, E., et al. (2021). Optogenetic Induction of Orbitostriatal Long-Term Potentiation in the Dorsomedial Striatum Elicits a Persistent Reduction of Alcohol- Seeking Behavior in Rats. Neuropharmacology 191, 108560.

Choi, S., and Lovinger, D.M. (1997). Decreased frequency but not amplitude of quantal synaptic responses associated with expression of corticostriatal long-term depression. J Neurosci 17, 8613–8620.

Criado, A.B., and Gomez e Segura, I.A. (2003). Reduction of isoflurane MAC by fentanyl or remifentanil in rats. Vet Anaesth Analg 30, 250–256.

Crittenden, J.R., and Graybiel, A.M. (2011). Basal Ganglia disorders associated with imbalances in the striatal striosome and matrix compartments. Front Neuroanat 5, 59.

Crittenden, J.R., and Graybiel, A.M. (2016). Disease-Associated Changes in the Striosome and Matrix Compartments of the Dorsal Striatum. Hbk Behav Neurosci 24, 783–802.

Crittenden, J.R., Tillberg, P.W., Riad, M.H., Shima, Y., Gerfen, C.R., Curry, J., Housman, D.E., Nelson, S.B., Boyden, E.S., and Graybiel, A.M. (2016). Striosome-dendron bouquets highlight a unique striatonigral circuit targeting dopamine-containing neurons. Proc Natl Acad Sci U S A 113, 11318–11323.

Cruikshank, S.J., Urabe, H., Nurmikko, A.V., and Connors, B.W. (2010). Pathway-specific feedforward circuits between thalamus and neocortex revealed by selective optical stimulation of axons. Neuron 65, 230–245.

Cunningham, C.L., Gremel, C.M., and Groblewski, P.A. (2006). Drug-induced conditioned place preference and aversion in mice. Nat Protoc 1, 1662–1670.

DeNardo, L.A., Liu, C.D., Allen, W.E., Adams, E.L., Friedmann, D., Fu, L., Guenthner, C.J., Tessier-Lavigne, M., and Luo, L. (2019). Temporal evolution of cortical ensembles promoting remote memory retrieval. Nat Neurosci 22, 460–469.

Deniau, J.M., Hammond, C., Chevalier, G., and Feger, J. (1978). Evidence for branched subthalamic nucleus projections to substantia nigra, entopeduncular nucleus and globus pallidus. Neurosci Lett 9, 117–121.

Di Chiara, G., and Imperato, A. (1988). Drugs abused by humans preferentially increase synaptic dopamine concentrations in the mesolimbic system of freely moving rats. Proc Natl Acad Sci U S A 85, 5274–5278.

Di Chiara, G., and North, R.A. (1992). Neurobiology of opiate abuse. Trends Pharmacol Sci 13, 185–193.

Diana, M. (2011). The dopamine hypothesis of drug addiction and its potential therapeutic value. Front Psychiatry 2, 64.

Dobrunz, L.E., and Stevens, C.F. (1997). Heterogeneity of release probability, facilitation, and depletion at central synapses. Neuron 18, 995–1008.

Drake, C.T., Aicher, S.A., Montalmant, F.L., and Milner, T.A. (2005). Redistribution of mu- opioid receptors in C1 adrenergic neurons following chronic administration of morphine. Exp Neurol 196, 365–372.

Evans, C.J. (2004). Secrets of the opium poppy revealed. Neuropharmacology 47 Suppl 1, 293–299.

Evans, R.C., Twedell, E.L., Zhu, M., Ascencio, J., Zhang, R., and Khaliq, Z.M. (2020). Functional Dissection of Basal Ganglia Inhibitory Inputs onto Substantia Nigra Dopaminergic Neurons. Cell Rep 32, 108156.

Ezeomah, C., Cunningham, K.A., Stutz, S.J., Fox, R.G., Bukreyeva, N., Dineley, K.T., Paessler, S., and Cisneros, I.E. (2020). Fentanyl self-administration impacts brain immune responses in male Sprague-Dawley rats. Brain Behav Immun 87, 725–738.

Faget, L., Zell, V., Souter, E., McPherson, A., Ressler, R., Gutierrez-Reed, N., Yoo, J.H., Dulcis, D., and Hnasko, T.S. (2018). Opponent control of behavioral reinforcement by inhibitory and excitatory projections from the ventral pallidum. Nature communications 9, 849.

Fricker, L.D., Margolis, E.B., Gomes, I., and Devi, L.A. (2020). Five Decades of Research on Opioid Peptides: Current Knowledge and Unanswered Questions. Molecular Pharmacology 98, 96–108.

Friedman, A., Homma, D., Bloem, B., Gibb, L.G., Amemori, K.I., Hu, D., Delcasso, S., Truong, T.F., Yang, J., Hood, A.S., et al. (2017). Chronic Stress Alters Striosome- Circuit Dynamics, Leading to Aberrant Decision-Making. Cell 171, 1191–1205 e1128.

Friedman, A., Homma, D., Gibb, L.G., Amemori, K., Rubin, S.J., Hood, A.S., Riad, M.H., and Graybiel, A.M. (2015). A Corticostriatal Path Targeting Striosomes Controls Decision-Making under Conflict. Cell 161, 1320–1333.

Friedman, A., Hueske, E., Drammis, S.M., Toro Arana, S.E., Nelson, E.D., Carter, C.W., Delcasso, S., Rodriguez, R.X., Lutwak, H., DiMarco, K.S., et al. (2020). Striosomes Mediate Value-Based Learning Vulnerable in Age and a Huntington’s Disease Model. Cell 183, 918–934 e949.

Fujiyama, F., Sohn, J., Nakano, T., Furuta, T., Nakamura, K.C., Matsuda, W., and Kaneko, T. (2011). Exclusive and common targets of neostriatofugal projections of rat striosome neurons: a single neuron-tracing study using a viral vector. Eur J Neurosci 33, 668–677.

Fujiyama, F., Takahashi, S., and Karube, F. (2015). Morphological elucidation of basal ganglia circuits contributing reward prediction. Frontiers in neuroscience 9, 6.

Fyfe, L.W., Cleary, D.R., Macey, T.A., Morgan, M.M., and Ingram, S.L. (2010). Tolerance to the antinociceptive effect of morphine in the absence of short-term presynaptic desensitization in rat periaqueductal gray neurons. J Pharmacol Exp Ther 335, 674–680.

Galaj, E., Han, X., Shen, H., Jordan, C.J., He, Y., Humburg, B., Bi, G.H., and Xi, Z.X. (2020). Dissecting the Role of GABA Neurons in the VTA versus SNr in Opioid Reward. J Neurosci 40, 8853–8869.

Gaulden, A.D., Burson, N., Sadik, N., Ghosh, I., Khan, S.J., Brummelte, S., Kallakuri, S., and Perrine, S.A. (2021). Effects of fentanyl on acute locomotor activity, behavioral sensitization, and contextual reward in female and male rats. Drug Alcohol Depend 229, 109101.

Gaveriaux-Ruff, C., and Kieffer, B.L. (2002). Opioid receptor genes inactivated in mice: the highlights. Neuropeptides 36, 62–71.

George, S.R., Zastawny, R.L., Brionesurbina, R., Cheng, R., Nguyen, T., Heiber, M., Kouvelas, A., Chan, A.S., and Odowd, B.F. (1994). Distinct Distributions of Mu, Delta and Kappa Opioid Receptor Messenger-Rna in Rat-Brain. Biochem Biophys Res Commun 205, 1438–1444.

Georges, F., Normand, E., Bloch, B., and Le Moine, C. (1998). Opioid receptor gene expression in the rat brain during ontogeny, with special reference to the mesostriatal system: an in situ hybridization study. Brain Res Dev Brain Res 109, 187–199.

Gerfen, C.R. (1984). The neostriatal mosaic: compartmentalization of corticostriatal input and striatonigral output systems. Nature 311, 461–464.

Gerfen, C.R., and Surmeier, D.J. (2011). Modulation of striatal projection systems by dopamine. Annu Rev Neurosci 34, 441–466.

Gopalakrishnan, L., Chatterjee, O., Ravishankar, N., Suresh, S., Raju, R., Mahadevan, A., and Prasad, T.S.K. (2021). Opioid receptors signaling network. J Cell Commun Signal.

Graybiel, A.M., and Ragsdale, C.W., Jr. (1978). Histochemically distinct compartments in the striatum of human, monkeys, and cat demonstrated by acetylthiocholinesterase staining. Proc Natl Acad Sci U S A 75, 5723–5726.

Grimm, J.W., Hope, B.T., Wise, R.A., and Shaham, Y. (2001). Neuroadaptation. Incubation of cocaine craving after withdrawal. Nature 412, 141–142.

Haberstock-Debic, H., Kim, K.A., Yu, Y.J., and von Zastrow, M. (2005). Morphine promotes rapid, arrestin-dependent endocytosis of mu-opioid receptors in striatal neurons. J Neurosci 25, 7847–7857.

Haberstock-Debic, H., Wein, M., Barrot, M., Colago, E.E., Rahman, Z., Neve, R.L., Pickel, V.M., Nestler, E.J., von Zastrow, M., and Svingos, A.L. (2003). Morphine acutely regulates opioid receptor trafficking selectively in dendrites of nucleus accumbens neurons. J Neurosci 23, 4324–4332.

Hack, S.P., Vaughan, C.W., and Christie, M.J. (2003). Modulation of GABA release during morphine withdrawal in midbrain neurons in vitro. Neuropharmacology 45, 575–584.

Hall, H., Sedvall, G., Magnusson, O., Kopp, J., Halldin, C., and Farde, L. (1994). Distribution of D1- and D2-dopamine receptors, and dopamine and its metabolites in the human brain. Neuropsychopharmacology 11, 245–256.

Hellard, E.R., Binette, A., Zhuang, X., Lu, J., Ma, T., Jones, B., Williams, E., Jayavelu, S., and Wang, J. (2019). Optogenetic control of alcohol-seeking behavior via the dorsomedial striatal circuit. Neuropharmacology 155, 89–97.

Heusler, P., Tardif, S., and Cussac, D. (2016). Agonist stimulation at human mu opioid receptors in a [S-35]GTP gamma S incorporation assay: observation of “bell- shaped” concentration-response relationships under conditions of strong receptor G protein coupling. J Recept Sig Transd 36, 158–166.

Hnasko, T.S., Sotak, B.N., and Palmiter, R.D. (2005). Morphine reward in dopamine- deficient mice. Nature 438, 854–857.

Hong, S., and Hikosaka, O. (2008). The globus pallidus sends reward-related signals to the lateral habenula. Neuron 60, 720–729.

Huang, C.C.Y., Ma, T., Roltsch Hellard, E.A., Wang, X., Selvamani, A., Lu, J., Sohrabji, F., and Wang, J. (2017). Stroke triggers nigrostriatal plasticity and increases alcohol consumption in rats. Sci Rep 7, 2501.

Jiang, Z.G., and North, R.A. (1992). Pre- and postsynaptic inhibition by opioids in rat striatum. J Neurosci 12, 356–361.

Johnson, S.W., and North, R.A. (1992). Opioids excite dopamine neurons by hyperpolarization of local interneurons. J Neurosci 12, 483–488.

Johnston, J.G., Gerfen, C.R., Haber, S.N., and van der Kooy, D. (1990). Mechanisms of striatal pattern formation: conservation of mammalian compartmentalization. Brain Res Dev Brain Res 57, 93–102.

Jordan, B.A., Cvejic, S., and Devi, L.A. (2000). Opioids and their complicated receptor complexes. Neuropsychopharmacology 23, S5–S18.

Koob, G.F. (2021). Drug addiction: hyperkatifeia/negative reinforcement as a framework for medications development. Pharmacol Rev 73, 163–201.

Koob, G.F., Powell, P., and White, A. (2020). Addiction as a Coping Response: Hyperkatifeia, Deaths of Despair, and COVID-19. Am J Psychiatry 177, 1031- 1037.

Koob, G.F., and Volkow, N.D. (2010). Neurocircuitry of addiction. Neuropsychopharmacology 35, 217–238.

Koob, G.F., and Volkow, N.D. (2016). Neurobiology of addiction: a neurocircuitry analysis. Lancet Psychiatry 3, 760–773.

Kovoor, A., Celver, J.P., Wu, A., and Chavkin, C. (1998). Agonist induced homologous desensitization of mu-opioid receptors mediated by G protein-coupled receptor kinases is dependent on agonist efficacy. Mol Pharmacol 54, 704–711.

Kravitz, A.V., Freeze, B.S., Parker, P.R., Kay, K., Thwin, M.T., Deisseroth, K., and Kreitzer, A.C. (2010). Regulation of parkinsonian motor behaviours by optogenetic control of basal ganglia circuitry. Nature 466, 622–626.

Kreitzer, A.C., and Malenka, R.C. (2008). Striatal plasticity and basal ganglia circuit function. Neuron 60, 543–554.

Lavian, H., Loewenstern, Y., Madar, R., Almog, M., Bar-Gad, I., Okun, E., and Korngreen, A. (2018). Dopamine receptors in the rat entopeduncular nucleus. Brain Struct Funct 223, 2673–2684.

Li, H., Eid, M., Pullmann, D., Chao, Y.S., Thomas, A.A., and Jhou, T.C. (2021a). Entopeduncular Nucleus Projections to the Lateral Habenula Contribute to Cocaine Avoidance. J Neurosci 41, 298–306.

Li, X., Rubio, F.J., Zeric, T., Bossert, J.M., Kambhampati, S., Cates, H.M., Kennedy, P.J., Liu, Q.R., Cimbro, R., Hope, B.T., et al. (2015). Incubation of Methamphetamine Craving Is Associated with Selective Increases in Expression of Bdnf and Trkb, Glutamate Receptors, and Epigenetic Enzymes in Cue-Activated Fos-Expressing Dorsal Striatal Neurons. J Neurosci 35, 8232–8244.

Li, Y., Simmler, L.D., Van Zessen, R., Flakowski, J., Wan, J.X., Deng, F., Li, Y.L., Nautiyal, K.M., Pascoli, V., and Luscher, C. (2021b). Synaptic mechanism underlying serotonin modulation of transition to cocaine addiction. Science 373, 1252–1256.

Li, Y.Q., Li, F.Q., Wang, X.Y., Wu, P., Zhao, M., Xu, C.M., Shaham, Y., and Lu, L. (2008).Central Amygdala Extracellular Signal-Regulated Kinase Signaling Pathway Is Critical to Incubation of Opiate Craving. J Neurosci 28, 13248–13257.

Lopez-Huerta, V.G., Nakano, Y., Bausenwein, J., Jaidar, O., Lazarus, M., Cherassse, Y., Garcia-Munoz, M., and Arbuthnott, G. (2016). The neostriatum: two entities, one structure? Brain Struct Funct 221, 1737–1749.

Lu, J., Cheng, Y., Wang, X., Woodson, K., Kemper, C., Disney, E., and Wang, J. (2019). Alcohol intake enhances glutamatergic transmission from D2 receptor-expressing afferents onto D1 receptor-expressing medium spiny neurons in the dorsomedial striatum. Neuropsychopharmacology 44, 1123–1131.

Lu, J., Cheng, Y., Xie, X., Woodson, K., Bonifacio, J., Disney, E., Barbee, B., Wang, X., Zaidi, M., and Wang, J. (2021a). Whole-brain mapping of direct inputs to dopamine D1 and D2 receptor-expressing medium spiny neurons in the posterior dorsomedial striatum. eNeuro 8, 0348–0320.2020.

Lu, J., Cheng, Y., Xie, X., Woodson, K., Bonifacio, J., Disney, E., Barbee, B., Wang, X., Zaidi, M., and Wang, J. (2021b). Whole-Brain Mapping of Direct Inputs to Dopamine D1 and D2 Receptor-Expressing Medium Spiny Neurons in the Posterior Dorsomedial Striatum. eNeuro 8.

Luscher, C., and Malenka, R.C. (2011). Drug-Evoked Synaptic Plasticity in Addiction: From Molecular Changes to Circuit Remodeling. Neuron 69, 650–663.

Luscher, C., and Ungless, M.A. (2006). The mechanistic classification of addictive drugs. PLoS Med 3, e437.

Ma, T., Cheng, Y., Roltsch Hellard, E., Wang, X., Lu, J., Gao, X., Huang, C.C.Y., Wei, X., Ji, J., and Wang, J. (2018). Bidirectional and long-lasting control of alcohol-seeking behavior by corticostriatal LTP and LTD. Nat Neurosci 21, 373–383.

Ma, T., Huang, Z., Xie, X., Cheng, Y., Zhuang, X., Childs, M.J., Gangal, H., Wang, X., Smith, L.N., Smith, R.J., et al. (2021). Chronic alcohol drinking persistently suppresses thalamostriatal excitation of cholinergic neurons to impair cognitive flexibility. J Clin Invest 132, e154969.

Madhavan, A., Bonci, A., and Whistler, J.L. (2010). Opioid-Induced GABA potentiation after chronic morphine attenuates the rewarding effects of opioids in the ventral tegmental area. J Neurosci 30, 14029–14035.

Maia, T.V., and Frank, M.J. (2011). From reinforcement learning models to psychiatric and neurological disorders. Nat Neurosci 14, 154–162.

Mansour, A., Meador-Woodruff, J.H., Zhou, Q., Civelli, O., Akil, H., and Watson, S.J. (1992). A comparison of D1 receptor binding and mRNA in rat brain using receptor autoradiographic and in situ hybridization techniques. Neuroscience 46, 959–971.

Matsui, A., Jarvie, B.C., Robinson, B.G., Hentges, S.T., and Williams, J.T. (2014). Separate GABA afferents to dopamine neurons mediate acute action of opioids, development of tolerance, and expression of withdrawal. Neuron 82, 1346–1356.

Matthes, H.W., Maldonado, R., Simonin, F., Valverde, O., Slowe, S., Kitchen, I., Befort, K., Dierich, A., Le Meur, M., Dolle, P., et al. (1996). Loss of morphine-induced analgesia, reward effect and withdrawal symptoms in mice lacking the mu-opioid- receptor gene. Nature 383, 819–823.

McGregor, M.M., McKinsey, G.L., Girasole, A.E., Bair-Marshall, C.J., Rubenstein, J.L.R., and Nelson, A.B. (2019). Functionally Distinct Connectivity of Developmentally Targeted Striosome Neurons. Cell Rep 29, 1419–1428 e1415.

Meye, F.J., Soiza-Reilly, M., Smit, T., Diana, M.A., Schwarz, M.K., and Mameli, M. (2016). Shifted pallidal co-release of GABA and glutamate in habenula drives cocaine withdrawal and relapse. Nat Neurosci 19, 1019–1024.

Milton, A.L., and Everitt, B.J. (2012). The persistence of maladaptive memory: addiction, drug memories and anti-relapse treatments. Neurosci Biobehav Rev 36, 1119–1139.

Missale, C., Nash, S.R., Robinson, S.W., Jaber, M., and Caron, M.G. (1998). Dopamine receptors: from structure to function. Physiol Rev 78, 189–225.

Miura, M., Masuda, M., and Aosaki, T. (2008). Roles of mu-opioid receptors in GABAergic synaptic transmission in the striosome and matrix compartments of the striatum. Mol Neurobiol 37, 104–115.

Miura, M., Saino-Saito, S., Masuda, M., Kobayashi, K., and Aosaki, T. (2007). Compartment-specific modulation of GABAergic synaptic transmission by mu-opioid receptor in the mouse striatum with green fluorescent protein-expressing dopamine islands. J Neurosci 27, 9721–9728.

Mizumori, S.J.Y., and Baker, P.M. (2017). The Lateral Habenula and Adaptive Behaviors. Trends Neurosci 40, 481–493.

Mollick, J.A., Hazy, T.E., Krueger, K.A., Nair, A., Mackie, P., Herd, S.A., and O’Reilly, R.C. (2020). A systems-neuroscience model of phasic dopamine. Psychol Rev 127, 972–1021.

Muelbl, M.J., Nawarawong, N.N., Clancy, P.T., Nettesheim, C.E., Lim, Y.W., and Olsen, C.M. (2016). Responses to drugs of abuse and non-drug rewards in leptin deficient ob/ob mice. Psychopharmacology (Berl) 233, 2799–2811.

Nadel, J.A., Pawelko, S.S., Scott, J.R., McLaughlin, R., Fox, M., Ghanem, M., van der Merwe, R., Hollon, N.G., Ramsson, E.S., and Howard, C.D. (2021). Optogenetic stimulation of striatal patches modifies habit formation and inhibits dopamine release. Sci Rep-Uk 11.

Petreanu, L., Mao, T., Sternson, S.M., and Svoboda, K. (2009). The subcellular organization of neocortical excitatory connections. Nature 457, 1142–1145.

Pickens, C.L., Airavaara, M., Theberge, F., Fanous, S., Hope, B.T., and Shaham, Y. (2011). Neurobiology of the incubation of drug craving. Trends Neurosci 34, 411–420.

Pothos, E., Rada, P., Mark, G.P., and Hoebel, B.G. (1991). Dopamine microdialysis in the nucleus accumbens during acute and chronic morphine, naloxone-precipitated withdrawal and clonidine treatment. Brain Res 566, 348–350.

Prager, E.M., and Plotkin, J.L. (2019). Compartmental function and modulation of the striatum. J Neurosci Res 97, 1503–1514.

Reiner, D.J., Fredriksson, I., Lofaro, O.M., Bossert, J.M., and Shaham, Y. (2019). Relapse to opioid seeking in rat models: behavior, pharmacology and circuits. Neuropsychopharmacology 44, 465–477.

Reuben, D.B., Alvanzo, A.A., Ashikaga, T., Bogat, G.A., Callahan, C.M., Ruffing, V., and Steffens, D.C. (2015). National Institutes of Health Pathways to Prevention Workshop: the role of opioids in the treatment of chronic pain. Ann Intern Med 162, 295–300.

Salinas, A.G., Mateo, Y., Carlson, V.C.C., Stinnett, G.S., Luo, G.X., Seasholtz, A.F., Grant, K.A., and Lovinger, D.M. (2021). Long-term alcohol consumption alters dorsal striatal dopamine release and regulation by D2 dopamine receptors in rhesus macaques. Neuropsychopharmacology 46, 1432–1441.

Samejima, K., Ueda, Y., Doya, K., and Kimura, M. (2005). Representation of action- specific reward values in the striatum. Science 310, 1337–1340.

Shabel, S.J., Proulx, C.D., Piriz, J., and Malinow, R. (2014). Mood regulation. GABA/glutamate co-release controls habenula output and is modified by antidepressant treatment. Science 345, 1494–1498.

Sim-Selley, L.J., Selley, D.E., Vogt, L.J., Childers, S.R., and Martin, T.J. (2000). Chronic heroin self-administration desensitizes mu opioid receptor-activated G-proteins in specific regions of rat brain. J Neurosci 20, 4555–4562.

Smith, J.B., Klug, J.R., Ross, D.L., Howard, C.D., Hollon, N.G., Ko, V.I., Hoffman, H., Callaway, E.M., Gerfen, C.R., and Jin, X. (2016a). Genetic-Based Dissection Unveils the Inputs and Outputs of Striatal Patch and Matrix Compartments. Neuron 91, 1069–1084.

Smith, J.B., Klug, J.R., Ross, D.L., Howard, C.D., Hollon, N.G., Ko, V.I., Hoffman, H., Callaway, E.M., Gerfen, C.R., and Jin, X. (2016b). Genetic-based dissection unveils the inputs and outputs of striatal patch and matrix compartments. Neuron 91, 1069–1084.

Stephenson-Jones, M., Kardamakis, A.A., Robertson, B., and Grillner, S. (2013). Independent circuits in the basal ganglia for the evaluation and selection of actions. Proc Natl Acad Sci U S A 110, E3670–3679.

Stephenson-Jones, M., Yu, K., Ahrens, S., Tucciarone, J.M., van Huijstee, A.N., Mejia, L.A., Penzo, M.A., Tai, L.H., Wilbrecht, L., and Li, B. (2016). A basal ganglia circuit for evaluating action outcomes. Nature 539, 289–293.

Stevenson, G.W., Giuvelis, D., Cormier, J., Cone, K., Atherton, P., Krivitsky, R., Warner, E., St Laurent, B., Dutra, J., Bidlack, J.M., et al. (2020). Behavioral pharmacology of the mixed-action delta-selective opioid receptor agonist BBI-11008: studies on acute, inflammatory and neuropathic pain, respiration, and drug self- administration. Psychopharmacology (Berl) 237, 1195–1208.

Sustkova-Fiserova, M., Puskina, N., Havlickova, T., Lapka, M., Syslova, K., Pohorala, V., and Charalambous, C. (2020). Ghrelin receptor antagonism of fentanyl-induced conditioned place preference, intravenous self-administration, and dopamine release in the nucleus accumbens in rats. Addict Biol 25, e12845.

Tan, T., Wang, W., Liu, T., Zhong, P., Conrow-Graham, M., Tian, X., and Yan, Z. (2021). Neural circuits and activity dynamics underlying sex-specific effects of chronic social isolation stress. Cell Rep 34, 108874.

Terman, G.W., Jin, W., Cheong, Y.P., Lowe, J., Caron, M.G., Lefkowitz, R.J., and Chavkin, C. (2004). G-protein receptor kinase 3 (GRK3) influences opioid analgesic tolerance but not opioid withdrawal. Br J Pharmacol 141, 55–64.

Thomsen, M., Woldbye, D.P., Wortwein, G., Fink-Jensen, A., Wess, J., and Caine, S.B. (2005). Reduced cocaine self-administration in muscarinic M5 acetylcholine receptor-deficient mice. J Neurosci 25, 8141–8149.

Tritsch, N.X., and Sabatini, B.L. (2012). Dopaminergic modulation of synaptic transmission in cortex and striatum. Neuron 76, 33–50.

Turchan, J., Przewlocka, B., Toth, G., Lason, W., Borsodi, A., and Przewlocki, R. (1999). The effect of repeated administration of morphine, cocaine and ethanol on mu and delta opioid receptor density in the nucleus accumbens and striatum of the rat. Neuroscience 91, 971–977.

Van den Oever, M.C., Goriounova, N.A., Li, K.W., Van der Schors, R.C., Binnekade, R., Schoffelmeer, A.N., Mansvelder, H.D., Smit, A.B., Spijker, S., and De Vries, T.J. (2008). Prefrontal cortex AMPA receptor plasticity is crucial for cue-induced relapse to heroin-seeking. Nat Neurosci 11, 1053–1058.

Varshneya, N.B., Walentiny, D.M., Moisa, L.T., Walker, T.D., Akinfiresoye, L.R., and Beardsley, P.M. (2021). Fentanyl-related substances elicit antinociception and hyperlocomotion in mice via opioid receptors. Pharmacology, biochemistry, and behavior 208, 173242.

Virk, M.S., and Williams, J.T. (2008). Agonist-specific regulation of mu-opioid receptor desensitization and recovery from desensitization. Mol Pharmacol 73, 1301–1308.

Volkow, N.D., Jones, E.B., Einstein, E.B., and Wargo, E.M. (2019). Prevention and Treatment of Opioid Misuse and Addiction A Review. Jama Psychiat 76, 208–216.

Wallace, M.L., Saunders, A., Huang, K.W., Philson, A.C., Goldman, M., Macosko, E.Z., McCarroll, S.A., and Sabatini, B.L. (2017). Genetically distinct parallel pathways in the entopeduncular nucleus for limbic and sensorimotor output of the basal ganglia. Neuron 94, 138–152 e135.

Watabe-Uchida, M., Zhu, L., Ogawa, S.K., Vamanrao, A., and Uchida, N. (2012). Whole- brain mapping of direct inputs to midbrain dopamine neurons. Neuron 74, 858–873.

Wei, X., Ma, T., Cheng, Y., Huang, C.C.Y., Wang, X., Lu, J., and Wang, J. (2018). Dopamine D1 or D2 receptor-expressing neurons in the central nervous system. Addiction biology 23, 569–584.

Williams, J.T., Christie, M.J., and Manzoni, O. (2001). Cellular and synaptic adaptations mediating opioid dependence. Physiol Rev 81, 299–343.

Williams, J.T., Ingram, S.L., Henderson, G., Chavkin, C., von Zastrow, M., Schulz, S., Koch, T., Evans, C.J., and Christie, M.J. (2013). Regulation of mu-opioid receptors: desensitization, phosphorylation, internalization, and tolerance. Pharmacol Rev 65, 223–254.

Xiao, X., Deng, H., Furlan, A., Yang, T., Zhang, X., Hwang, G.R., Tucciarone, J., Wu, P., He, M., Palaniswamy, R., et al. (2020). A Genetically Defined Compartmentalized Striatal Direct Pathway for Negative Reinforcement. Cell 183, 211–227 e220.

Yao, J., Zhang, Q., Liao, X., Li, Q., Liang, S., Li, X., Zhang, Y., Li, X., Wang, H., Qin, H., et al. (2018). A corticopontine circuit for initiation of urination. Nat Neurosci 21, 1541–1550.

Yu, Y.J., Arttamangkul, S., Evans, C.J., Williams, J.T., and von Zastrow, M. (2009). Neurokinin 1 receptors regulate morphine-induced endocytosis and desensitization of mu-opioid receptors in CNS neurons. J Neurosci 29, 222–233.

Zhang, G.W., Shen, L., Zhong, W., Xiong, Y., Zhang, L.I., and Tao, H.W. (2018). Transforming Sensory Cues into Aversive Emotion via Septal-Habenular Pathway. Neuron 99, 1016–1028 e1015.

Zingg, B., Chou, X.L., Zhang, Z.G., Mesik, L., Liang, F., Tao, H.W., and Zhang, L.I. (2017). AAV-mediated anterograde transsynaptic tagging: Mapping corticocollicular input- defined neural pathways for defense behaviors. Neuron 93, 33–47.

Zingg, B., Peng, B., Huang, J., Tao, H.W., and Zhang, L.I. (2020). Synaptic Specificity and Application of Anterograde Trans-Synaptic AAV for Probing Neural Circuitry. J Neurosci 4, 3250–3267.

Zucker, R.S., and Regehr, W.G. (2002). Short-term synaptic plasticity. Annu Rev Physiol 64, 355–405.

